# Transcranial electrical stimulation for memory enhancement: A systematic review and meta-analysis

**DOI:** 10.1101/2025.03.17.643623

**Authors:** Yujin Goto, Ryoji Onagawa, Mitsuaki Takemi, Keiichi Ishikawa, Suzuka Narukawa, Kaoru Amano, Koichi Hosomi

## Abstract

Non-invasive brain stimulation techniques have received increasing interest for their potential to enhance memory function, a fundamental cognitive aspect of daily life.

This systematic review and meta-analysis aimed to investigate the efficacy of transcranial electrical stimulation (tES) in enhancing memory function in healthy adults, following the pre-registered strategy at PROSPERO (CRD42022353630).

A total of 113 articles (193 trials, 7,299 participants) examining tES were identified. Meta-analysis revealed a significant overall effect of tES on memory function compared with sham stimulation (standardised mean difference [95% confidence interval] = 0.19 [0.12, 0.25]). A positive pooled effect was also observed in the subgroup receiving frontal transcranial direct current stimulation (tDCS). Stimulation was associated with positive pooled effects for working- and declarative-memory task performance. Significant pooled effects were observed within 1 day after stimulation; longer-term estimates were imprecise. Regarding adverse events, tingling and itching sensations on the scalp occurred more frequently in the active group than in the sham group, but no serious adverse events were reported. Limitations, including small-study effects, heterogeneity, and uneven coverage of memory domains, were noted, emphasising the need for improved experimental rigour and diversification of memory tasks.

These findings suggest that tES is associated with small improvements in memory-task performance and was generally well tolerated under the studied protocols. Heterogeneity, small-study effects, and limited adverse-event reporting constrain conclusions about efficacy, durability, and safety.

## 1. Introduction

Memory plays a crucial role in daily life, influencing both routine tasks and professional performance. In today’s information-saturated world, the acquisition and effective application of information are critical. Various approaches, including pharmacological interventions (Fond et al., 2015), hormonal treatments (Gold, 2008), and cognitive training (Morrison & Chein, 2011; Redick et al., 2013), have been explored for memory enhancement. However, challenges related to reliability, ethical considerations, and widespread implementation (Barch, 2004; Fond et al., 2015; Morrison & Chein, 2011; Redick et al., 2013) often accompany these approaches. Consequently, there is growing interest in non-invasive, safe, and cost-effective techniques that can be widely implemented. Among such methods, non-invasive brain stimulation (NIBS) has emerged as a promising and rapidly evolving technique.

Transcranial electrical stimulation (tES) is exceptional given its relatively simple design, making it accessible to both researchers and general users (Antal et al., 2017; Woods et al., 2015). In contrast, other NIBS techniques, such as transcranial magnetic stimulation (TMS) (Rossi et al., 2009) and transcranial focused ultrasound stimulation (tFUS) (Legon et al., 2014), require larger and more expensive equipment, limiting their widespread use. The most widely used forms of tES are transcranial direct current stimulation (tDCS) and transcranial alternating current stimulation (tACS). tDCS applies a weak constant current that can bias neuronal membrane potentials and alter activity-dependent plasticity, whereas tACS applies an oscillatory current that may interact with endogenous rhythms and modify the timing or synchronisation of neural activity (Kanai et al., 2008; Nitsche et al., 2008; Woods et al., 2015). tDCS montages vary in the number, size, polarity, and positions of the electrodes. In this review, montages were grouped as monopolar, bipolar, or high-definition (HD) tDCS for descriptive and subgroup analyses. Here, in bipolar montages, the return electrode is placed contralaterally to the target. In monopolar montages, the target electrode is positioned over the intended cortical region, with the return electrode placed at an extracephalic site to concentrate current around the target. High-definition (HD) tDCS enhances focality by surrounding the target electrode with multiple smaller return electrodes, thereby constraining the current distribution and minimising directional bias in the induced electric field. Other tES modalities, including transcranial random noise stimulation (tRNS) and transcranial pulsed current stimulation (tPCS), have also been investigated for their potential applications in NIBS (Woods et al., 2015).

However, the behavioural effects of either technique cannot be inferred from the stimulus type and electrode position only, since current flows between all electrodes, and the resulting electric field depends on the current intensity, waveform, electrode geometry, individual anatomy, brain state during the stimulation, and stimulation timing (Laakso et al., 2015; Li et al., 2015; Miniussi et al., 2013; Woods et al., 2015). This variability is especially relevant to memory research, in which stimulation has been applied to frontal, parietal, temporal, and distributed frontoparietal configurations, often during different stages of encoding, maintenance, consolidation, or retrieval.

Previous reviews have reported effects ranging from negligible to modest, with conclusions differing across memory domains, participant populations, stimulation techniques, and experimental designs (Barham et al., 2016; Brunoni & Vanderhasselt, 2014; Chaieb et al., 2008; Galli et al., 2018; Goldthorpe et al., 2020; Hill et al., 2015; Horvath et al., 2015; Laakso et al., 2015; Li et al., 2015; Mancuso et al., 2016; Tremblay et al., 2014; Woods et al., 2015). Several reviews have focused on specific outcomes, such as working memory, episodic memory, sleep-related consolidation, or healthy ageing, while others have pooled heterogeneous populations and protocols. Consequently, the overall magnitude and durability of tES effects in healthy adults remain uncertain, as do the contributions of stimulation modality, montage, age, memory domain, and assessment timing. The correspondence between behavioural effects and the induced cortical electric field is also limited. Furthermore, broader inclusion criteria in previous reviews may have included studies with questionable validity or reliability, potentially compromising overall conclusions. A more rigorous evaluation of tES for memory enhancement requires stricter methodological criteria alongside a balanced assessment of its potential benefits and risks to ensure a well-balanced perspective on its implications.

We therefore conducted a preregistered systematic review and multilevel meta-analysis of parallel-group randomised controlled trials examining tES and memory performance in healthy adults. First, we mapped the development and composition of the literature, including stimulation modalities and montages, memory domains, task characteristics, sensory modalities, and assessment timing. Second, we estimated the overall effect of tES and examined variation according to stimulation type, age, memory domain, timing, and stimulation site. Finally, we combined electric-field modelling with meta-analytic effect sizes to identify cortical regions in which stronger stimulation was associated with greater behavioural improvement. By integrating these, this review aimed to clarify not only whether tES alters memory-task performance, but also the conditions and neural distributions under which such effects are most consistently observed.

## 2. Methods

This review examined the question: “How does non-invasive brain stimulation influence memory function in healthy adults compared to control conditions?” To ensure methodological rigour, this systematic review and meta-analysis followed the Minds Handbook for Clinical Practice Guideline Development 2020 (version 3.0, chapter 4) (Minds Manual Developing Committee, 2021). The PRISMA checklist is shown in Supplementary Table 1. A pre-registration strategy (PROSPERO, CRD42022353630) was implemented prior to initiating the database search.

### 2.1. Database search

To address the review question, we selected studies that examined the effects of NIBS on memory function in healthy adults using an appropriate randomised controlled trial (RCT) design.

We conducted a comprehensive search across PubMed, Scopus, Web of Science, PsycINFO, JDreamIII, and Ichushi for peer-reviewed articles available in English or Japanese. These databases were selected based on a previous meta-analysis (Kimura et al., 2024) to maximise coverage. The search, conducted on January 29, 2026, focused on two key domains—“non-invasive brain stimulation” and “memory.” Detailed queries are available at https://osf.io/rfu3n/. Duplicate records were removed before screening; irrelevant articles were excluded during title and abstract screening.

### 2.2. Article selection and data extraction

To ensure a comprehensive systematic review and meta-analysis, we screened articles that met the following inclusion criteria: (1) participants were healthy adults, (2) the average participant age was over 18 years; (3) NIBS methods—excluding TMS, tFUS, and PNS— were used to enhance memory, as outlined in the introduction, given their limited suitability for general application; (4) the publication was an original peer-reviewed article or conference paper written in English or Japanese; (5) a parallel randomised controlled trial was conducted; (6) memory performance was assessed for working, episodic, associative, or semantic domains; and (7) memory performance data were sufficiently extractable for effect size calculation. (8) For inclusion in the meta-analysis, studies also needed to demonstrate a low or moderate risk of bias (see Section 2.3. Quality Assessment).

A two-phase screening process was conducted to identify relevant studies. Two reviewers independently performed each screening, and all six reviewers discussed studies where disagreements arose until a consensus was reached. The first screening phase involved eliminating irrelevant studies based on titles and abstracts. Studies that met consensus criteria advanced to the next phase. The second screening phase included a full-text assessment, ensuring non-adherence to the inclusion criteria was excluded at this stage.

During the second screening, each reviewer independently extracted key data, including study design, participant demographics, NIBS details, and memory outcome measures.

### 2.3. Quality assessment

Four authors assessed the risk of bias using the Cochrane Risk of Bias Assessment tool (Higgins et al., 2011), rating each item as “high”, “unclear”, or “low” risk. Items were grouped into six domains: selection bias (randomisation, baseline imbalance, allocation concealment), performance bias, detection bias, attrition bias (intention-to-treat analysis, incomplete outcome reporting), reporting bias, and other bias (early trial termination and other sources of bias, including conflicts of interest, inadequate sample size determination, and statistical method inconsistencies). A domain with several items was rated “high” if any item was “high”, “unclear” if any item was “unclear” and none was “high”, and “low” if all items were “low”. The overall rating followed an operational scoring rule of this review, which is not the overall-judgement algorithm of the Cochrane tool. A score was calculated as 2 × (number of “high” ratings) + (number of “unclear” ratings) across seven entries: the summary ratings of selection and attrition bias, the ratings of performance, detection, and reporting bias, and the two items of other bias. The overall risk was classified as high if the score was 8 or more, or if three or more “high” ratings were present among the item-level and summary ratings; as low if the score was 4 or less and no more than one “high” rating was present; and as “unclear” (intermediate) otherwise.

Four authors assessed the level of indirectness in all included studies. Indirectness was evaluated in four domains: population, intervention, comparison, and outcome, and it was categorised as “high”, “unclear”, or “low”. The overall level of indirectness was classified as “high” if three or more domains were rated “high”, moderate if two domains were rated “high”, and “low” in all other cases.

### 2.4. Data analyses

We conducted a standard pairwise comparison (active vs sham stimulation) using a multilevel meta-analysis with a random-effects model to evaluate the effectiveness of NIBS on memory performance. The analysis was conducted with the rma.mv function from the metafor package in R ver. 4.3.2. Individual effect sizes, expressed as standardised mean differences (SMDs), were assumed to be nested within articles, studies, and memory tasks. Effect sizes were Hedges’ g computed from the means, standard deviations, and sample sizes of the active and sham groups (esc package) and signed so that positive values indicate better memory performance under active stimulation; for seven effect sizes from three articles in which means and standard deviations could not be extracted, g was derived from reported t values. In rma.mv, the nesting was specified as random intercepts for article, study group within article, and task within study group (random = ∼ 1 | study ID/trial/memory task), and the sampling variance of each effect size was entered as a diagonal sampling-variance matrix. Dependence arising from comparisons that share a control group or the same participants was therefore accommodated only through these nested random effects; covariances between sampling errors were not modelled. To assess potential publication bias (systematic skew in published results), we used funnel plots along with Begg’s and Egger’s statistical tests, applying a significance threshold of *p* < 0.1, in accordance with Chapter 4 of the Minds Manual (Minds Manual Developing Committee, 2021). Egger’s test was implemented within the same multilevel model, with the standard error entered as a moderator and the test of its slope reported, so that dependent effect sizes were not treated as independent. Begg’s rank-correlation test has no multilevel counterpart and was applied to all effect sizes as if they were independent; it should therefore be regarded as approximate. Between-study variability, or heterogeneity, was tested using Cochran’s Q test.

We first assessed the overall effect of NIBS on memory and then conducted subgroup analyses to evaluate the effects of modulation across several categorical aspects. Specifically, the effects of two types of brain stimulation―tDCS and tACS―were tested. Other stimulation types (tRNS and combined tDCS–tACS or tDCS–tRNS protocols; 8 trials in total) contributed to the overall estimate but were not analysed as separate subgroups or included in the tDCS–tACS comparison, owing to the small number of related articles. Given that these stimulation modalities affect neuronal population activity via different mechanisms, analysing stimulation effects by stimulation type can help identify effective methodologies and provide insights into the underlying neural mechanisms (Miniussi et al., 2013).

Furthermore, subgroup analyses were performed for tDCS. The subgroup analysis included the following: (1) electric montage (monopolar, bipolar, HD); (2) age groups (participants younger than 65 years or aged 65 years and older); (3) type of memory function (working memory or declarative memory—including episodic, associative, and semantic memory); (4) timing of memory assessment relative to brain stimulation: during stimulation (online), within hours, one day later, one week later, or more than one week later to assess the durability of tES effects; and (5) stimulation of the frontal brain region (anode location at F3 or F4) using HD or monopolar tDCS, where a sufficient number of studies were available (Refer to section 3.2. Articles and Sample Characteristics). For each of these categorical subgroup dimensions, we additionally tested whether the levels differed significantly from one another using an omnibus test of moderators (QM statistic) within the corresponding multilevel meta-regression model.

We performed a meta-analysis using a random-effects model to determine whether adverse events were more likely with active stimulation than with sham stimulation. The negative effects of active stimulation were evaluated using SMDs for scale-based indicators, such as the visual analogue scale or numerical rating scale, and log-risk ratios for the number of adverse event cases. Trials in which no adverse events occurred in either the active or the sham group were excluded from the risk-ratio meta-analyses, as such trials provide no information about the relative risk between groups (Higgins et al., 2011). Subgroup analyses were performed for all data as well as for specific types of adverse events, including burning and skin redness, tingling and itching, fatigue, headache and neck pain, concentration problems, mood changes, discomfort, and sleepiness, and overall ratings (composite or global ratings of side effects, such as total or averaged side-effect scores or overall discomfort ratings; scale-based indicators only). Additionally, the proportion of participants reporting each type of adverse event was pooled separately for the active and the sham group using a random-effects model. These proportions were pooled on the logit scale and back-transformed using the logistic function, which constrains the resulting confidence intervals to lie between 0% and 100%. The double-zero exclusion described above was not applied to these per-arm proportions, because the observed incidence within a single arm remains informative about that arm’s event rate even when neither group reported an event.

### 2.5. Meta-analytic associations between electric field distribution and memory performance

To investigate the relationship between memory performance changes and tDCS-induced electric fields, we performed electric field modelling and regression analyses. Electric field simulations were performed using SimNIBS 4.5.0 (Thielscher et al., 2015) with the “Ernie extended” individualised head model (Van Hoornweder et al., 2024), using previously validated tissue conductivity values (Windhoff et al., 2013). Criteria for study inclusion required a consistent tDCS montage across experimental conditions and sufficient reporting of electrode position, shape, size, and material for reproduction. We interpolated electric field magnitudes to the middle depth of the cortical grey matter before resampling to the FsAverage surface. Cortical parcellation followed the HCP-MMP 1.0 atlas, comprising 360 regions of interest (ROIs; Glasser et al., 2016).

Building on the performance–electric field correlation (PEC) meta-analytic procedure (Wischnewski et al., 2021), we examined associations between Hedges’ g and two predictors: electric field strength (E) and electric field strength multiplied by cumulative stimulation duration in seconds (E × T). Based on the subgroup meta-analysis, assessments were restricted to the early testing intervals corresponding to timing categories 1–3: during, immediately after, or within 24 hours of tDCS. Separate analyses were conducted for working memory (WM) and declarative memory (DM), yielding four model-by-memory-type combinations. All age groups and montage types were retained.

When studies provided multiple effect sizes for the same combination of study, montage, memory type, age group, stimulation duration, and assessment timing, we calculated the weighted mean of Hedges’ g using the weights derived from the meta-analysis. The weights were the row-sum weights (weights(type = “rowsum”) in metafor) of the multilevel random-effects model fitted to the sham-controlled effect sizes of the tDCS articles eligible for field simulation, normalised to sum to 100. Observations with different stimulation durations or assessment timings were retained separately. For each ROI, E was calculated as the surface-area-weighted mean electric field magnitude. E × T was calculated after regional averaging. Separate vertex-wise models used the corresponding vertex-level predictor. We then fitted separate ordinary least-squares models for each predictor and ROI, adjusting for age group and memory test timing relative to tDCS as categorical covariates:

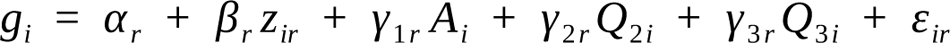

Here, i denotes an aggregated observation, r denotes an ROI, g is Hedges’ g, and z is the standardised E or E × T predictor. A indicates the older age group, with younger adults as the reference group. Q₂ and Q₃ indicate assessment-timing categories 2 and 3, with category 1 as the reference. Within each memory group and ROI, predictors were centred and divided by their standard deviation. The coefficient β therefore represents the change in Hedges’ g per one standard deviation increase in the predictor. Standard errors were clustered by study using the CR1 finite-sample correction to account for dependence among observations from the same study (Cameron & Miller, 2015). Each t-statistic was calculated as the predictor coefficient divided by its corresponding standard error. We applied one-sided tests for positive associations based on the directional hypothesis that greater memory enhancement would be associated with higher E or E × T.

ROI-level inference used 10,000 null-imposed wild cluster bootstrap samples (Cameron et al., 2008), with maximum-t (maxT) correction for multiple comparisons across 360 ROIs (Nichols & Holmes, 2002). We first fitted a reduced model containing the intercept and covariates but excluding the tested predictor. For each bootstrap sample, reduced-model residuals were multiplied by study-specific random signs (+1 or −1 with equal probability) and added to the reduced-model fitted values. The full model was then refitted, and t-statistics were recalculated using CR1 standard errors. The same sign was applied to all observations from a study, with identical sign assignments across ROIs and predictors within each memory group.

For each bootstrap sample, the largest signed t-statistic across the 360 ROIs was retained, yielding a null distribution of the maximum t-statistic. Each observed ROI t-statistic was compared with this distribution to obtain a family-wise error (FWE)-corrected p-value. Corrected p-values were calculated as (k + 1)/(B + 1), where k is the number of bootstrap maxima at least as large as the observed statistic and B = 10,000. Statistical significance was defined as one-sided FWE-corrected p < 0.05, separately for each predictor and memory group.

To assess the influence of individual studies, regression coefficients were re-estimated for all significant ROIs after excluding each study and all its observations. For the representative ROI with the smallest corrected p-value, individual bootstrap tests were also repeated using 10,000 samples per exclusion, with the ROI and full-sample predictor standardisation held fixed. These sensitivity p-values were calculated for the fixed ROI without repeating the 360-ROI correction. Analyses were implemented in Python 3.12.14.

## 3. Results

### 3.1. Search results

Figure 1 presents a flowchart of the review. We obtained 16,389 unique articles through database searches. In the initial screening phase, 15,817 irrelevant articles were excluded based on their titles and abstracts. As a result, 572 articles remained for the full-text eligibility assessment. Through full-text assessment, 459 articles were excluded for the following reasons: not being an original peer-reviewed article (n = 36), article retraction (n = 1), using the same data as another article (n = 3), non-RCT (n = 146), not a parallel design (n = 177), insufficient information regarding group comparison (n = 2), utilisation of TMS (n = 3), not the targeted population (n = 2), lack of memory-related outcome measures (n = 27), and unavailable data even after contacting the authors (n = 62). Finally, 113 articles were included in the qualitative analysis. Two articles having a high risk of bias, one article in which the effect size was a substantial outlier, and two articles that lacked a sham control were excluded before the meta-analysis and safety assessment.

**Figure 1.**
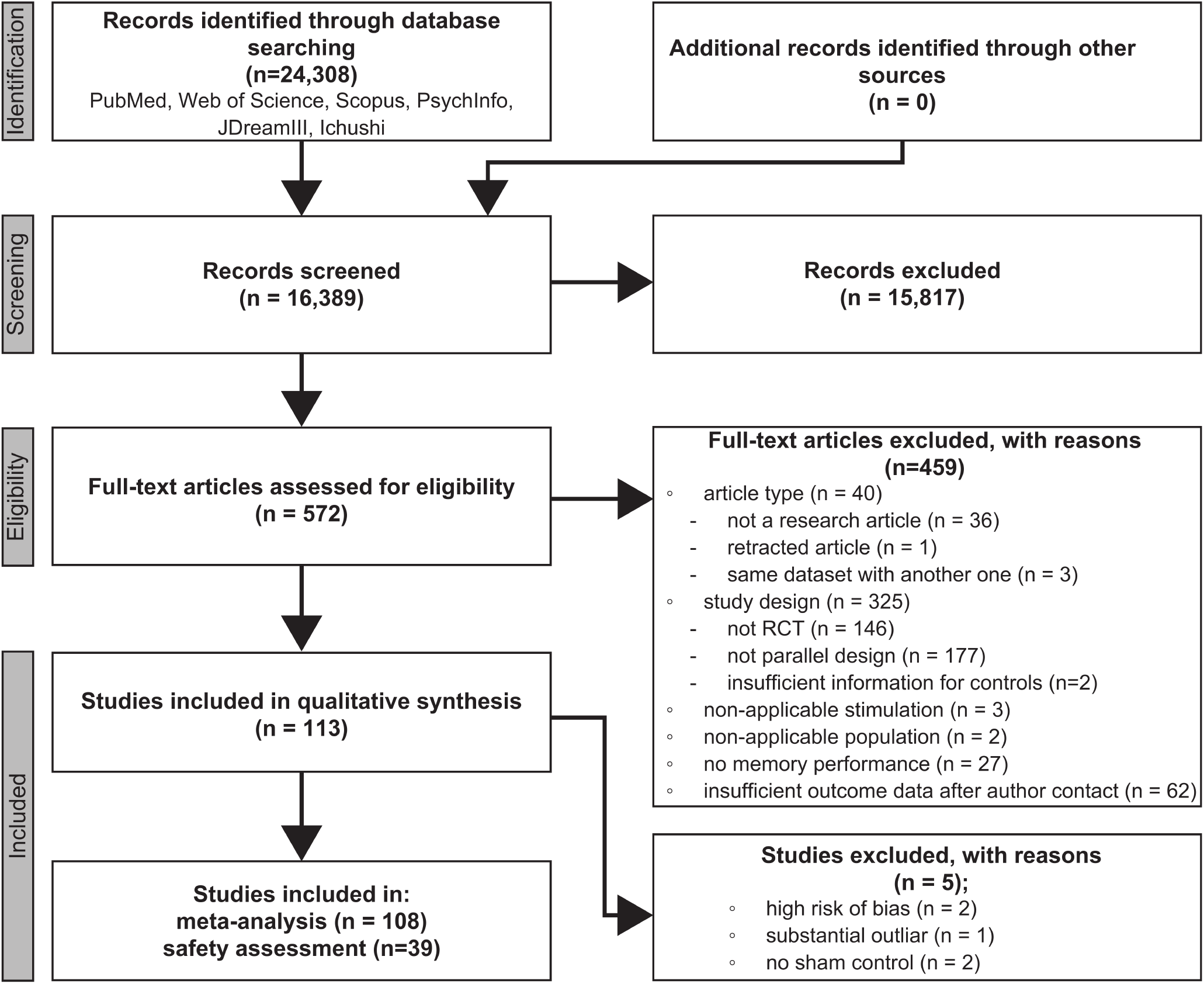
PRISMA flowchart of article search and screening.

### 3.2. Article and sample characteristics

Supplementary File 1 provides a summary of the 113 articles included in the qualitative analysis. The total number of participants was 7,299, with a mean age of 34.21 years (±20.72). The reported male-to-female ratio was 2,389 to 3,532. The participant numbers were extracted separately in cases where both total and sex-separated data were reported. A complete version of all extracted information is available at https://osf.io/83s94.

The publication years of articles that passed the first screening, along with their classification results, are shown in Fig. 2A. The earliest article that met the first screening criteria was published in 2004, and the first article to pass the second screening was published in 2013. Notably, just over half of the accepted articles (57 of 113; 50.4%) were published between 2022 and 2025. Authors of 86 studies lacking publicly available data were contacted, and 23 responded.

**Figure 2.**
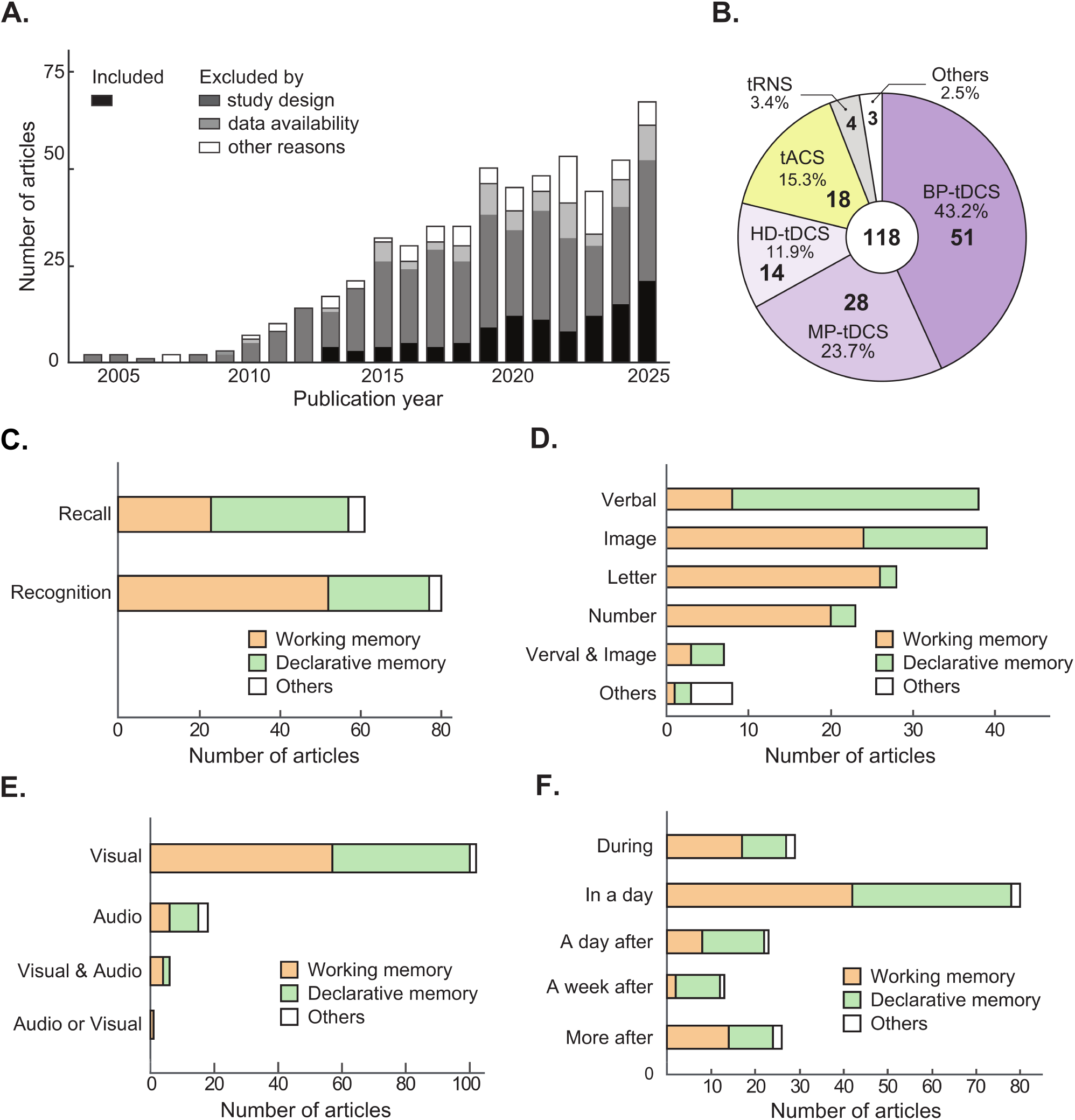
Summary of qualitative assessment. (A) Number of articles included or excluded at full-text screening, by publication year. Colours indicate inclusion or the reason for exclusion. (B) Stimulation methods and electrode-montage categories: bipolar tDCS (BP-tDCS), monopolar tDCS (MP-tDCS), high-definition tDCS (HD-tDCS), tACS, tRNS, and Others. All montage types are combined within the tACS and tRNS categories. Others comprises one other tDCS montage, one combined tDCS/tACS protocol, and one combined tDCS/tRNS protocol. Numbers and percentages in the sectors indicate the count and proportion of each category. The central number (118) is the sum of these category counts; articles reporting more than one stimulation or montage category contribute to each relevant category. (C) Memory-test types (recall or recognition). (D) Types of material used in the memory tasks. (E) Sensory modalities in which the memory-task stimuli were presented. (F) Timing of memory assessment relative to stimulation. “During” indicates assessment during stimulation; “In a day” indicates assessment within 24 h of the last stimulation session. The remaining categories indicate assessment one day, one week, or more than one week after stimulation. In C–F, bars show article counts and colours distinguish working memory, declarative memory, and other memory functions. Articles may contribute to more than one category. tDCS, transcranial direct current stimulation; tACS, transcranial alternating current stimulation; tRNS, transcranial random noise stimulation.

In the subsequent assessment of sample characteristics, articles that met multiple categories were counted separately. For example, an article with three independent experiments—Exp. 1: tDCS assessment for working memory, Exp. 2: tACS for working memory, and Exp. 3: tDCS for declarative memory—was counted once for each category: tDCS, tACS, working memory, and declarative memory. As a result, the summed counts across categories exceeded 113—the number of studies included in the qualitative synthesis, as shown in Fig. 1.

The most commonly used stimulus type and electrode montage was anodal tDCS targeting the frontal region (Fig. 2B, 3A–C). Ninety-four articles (83.2%) used tDCS, and eighteen articles (15.9%) used tACS. Four articles examined tRNS, and one article each examined a combination of tDCS and tACS and a combination of tDCS and tRNS. One tDCS article used an Other-tDCS montage, which is included in Others in Fig. 2B. Figures 3A–C show the placement of electrodes for tDCS stimulation of monopolar (Fig. 3A), bipolar (Fig. 3B, the full version of this figure is shown in Supplementary Figure 1), and HD-tDCS (Fig. 3C), and Figures 3D–F show the electrode montages of tACS in the same order. More than half of the articles targeted the frontal area (68 for tDCS, 12 for tACS), especially the DLPFC (58 for tDCS, 8 for tACS), with at least one electrode placed on F3 or F4 (62 for tDCS and 8 for tACS) following the international 10-10 EEG system. Accordingly, a subgroup labelled “stimulated frontal brain regions” in the subgroup analysis was created for further analysis.

**Figure 3.**
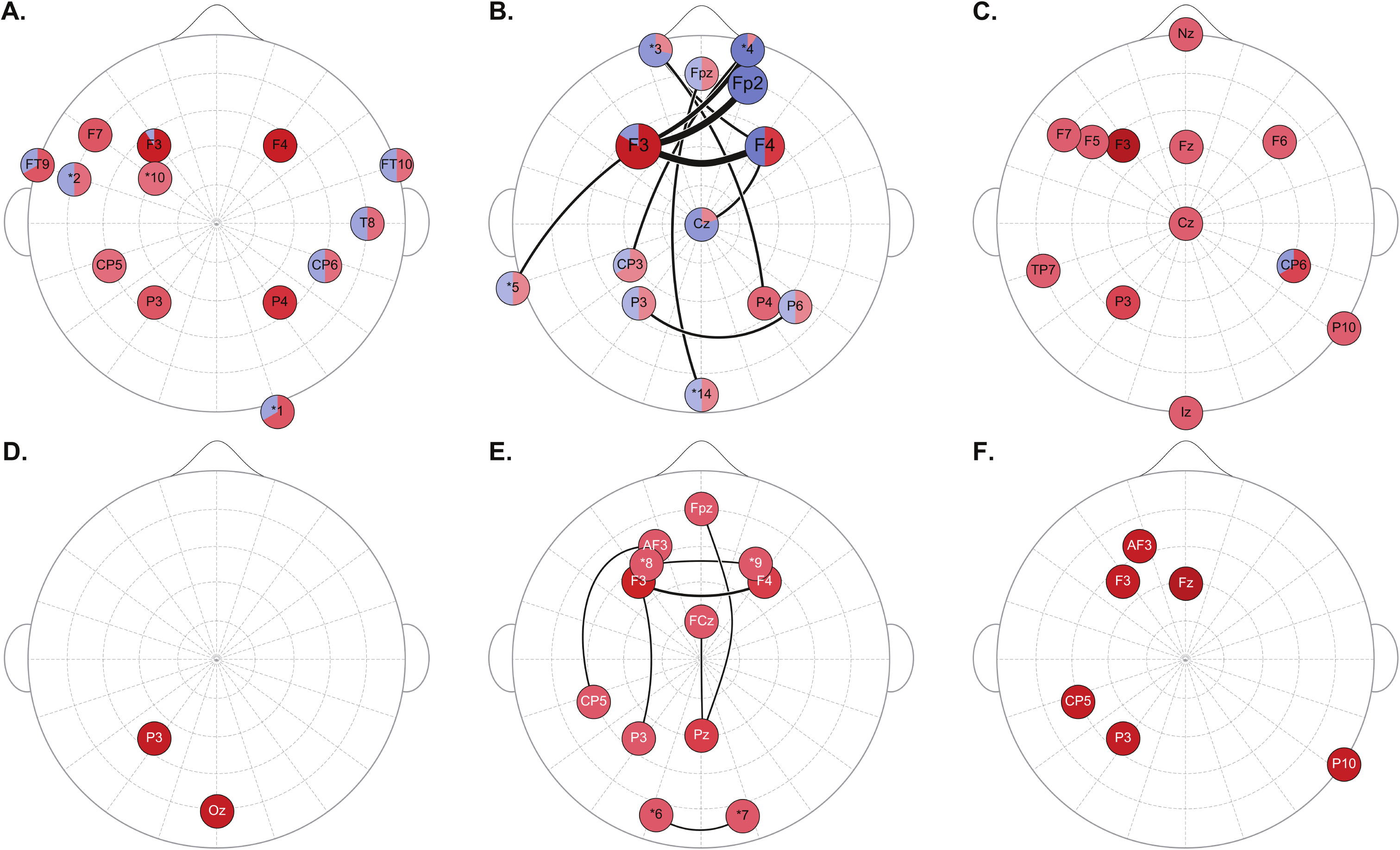
Electrode montages. Electrode montages for tDCS (A–C) and tACS (D–F), shown on the international 10–10 electrode layout: (A) monopolar tDCS, (B) bipolar tDCS, (C) high-definition (HD) tDCS, (D) monopolar tACS, (E) bipolar tACS, and (F) HD-tACS. Red and blue sectors in A–C indicate anodal and cathodal use, respectively; their proportions reflect the corresponding article counts summed across electrode configurations. An article reporting multiple configurations or polarities may contribute more than once. Colour intensity in A–C represents these counts in four bands (1, 2–3, 4–7, and ≥8). Node sizes in A –C are adjusted for display and do not quantitatively encode article counts. In D–F, red marks electrode locations and does not indicate a fixed polarity. Black curves connect paired electrodes, with thicker curves indicating larger article counts. For bipolar tDCS, counts for the two electrode directions are added; an article using both polarities can contribute to both directions. Panel B shows only pairs with identifiable electrode locations and a summed count of at least two, together with the endpoints of those pairs. The corresponding bipolar tDCS map, including pairs with a summed count of one, is shown in Supplementary Figure 1. Connections involving Fp3 are omitted because its scalp location could not be established; no coordinate was inferred. For monopolar and HD montages, only the main or centre target electrodes are shown. Multisite configurations were not decomposed into inferred pairwise connections. The chin electrode is shown schematically outside the scalp and does not represent an EEG coordinate. Asterisks denote nonstandard placements: *1, right cerebellum; *2, midpoint between F7, T7, and FT9; *3, left supraorbital area; *4, right supraorbital area; *5, left mastoid; *6, left C2 nerve dermatome; *7, right C2 nerve dermatome; *8, midpoint between F3 and AF3; *9, midpoint between F4 and AF4; *10, intersection of the F7–Cz and T7–Fz lines; and *14, midpoint between the inion (Iz) and Oz. T7 is the current designation for the former T3 position. tDCS, transcranial direct current stimulation; tACS, transcranial alternating current stimulation.

As for the memory type and task characteristics examined in the included studies, the evaluated articles focused more on working memory than declarative memory, with more on recognition-than recall-related memory (Fig. 2C–F). Specifically, 67 articles examined working memory, and 52 focused on declarative memory. Since declarative memory in this review includes episodic, semantic, and associative memory, working memory emerged as the most frequently investigated domain. Unlike recall tasks, which exhibited a few more declarative, but roughly balanced ratio of memory types, most of the articles on recognition tasks assessed working memory. This pattern is consistent with the use of n-back tasks, a well-known paradigm for evaluating working memory.

The types of memorised items varied widely but were generally simple, consisting of visually presented, low-information content such as single digits (Figs. 2D, E). Memory targets included verbal material (e.g., words), images, letters, and numbers. Six articles used audiovisual presentation, and one used visual or auditory presentation. Research focusing on verbal stimuli has dominantly examined declarative memory, whereas studies using other items have primarily investigated working memory (Fig. 2D). This discrepancy has resulted in an imbalanced ratio of memory types across different item categories. Visual presentation was the most common modality (92 of 113 studies; 81.4%), followed by auditory, audiovisual, or mixed formats.

Regarding the timing of memory assessment, 25.7% of studies evaluated performance online during stimulation, whereas 83.2% of studies assessed memory after stimulation to examine its after-effects (Fig. 2F). The most common approach, accounting for 64.6% of studies, was to test memory later on the same day. These assessments varied considerably in timing, including immediately after stimulation, following a short rest period, or after completion of an intervening task unrelated to memory. By contrast, relatively few studies assessed medium- or long-term effects beyond one day after stimulation.

### 3.3. Risk of bias assessment

The overall risk of bias was “high” in 2 studies, “unclear” in 66 studies, and “low” in 45 studies (Fig. 4A). In the domain of selection bias, 88 studies did not report the method used for randomising participants into groups. For performance bias, 71 studies were assessed as having a low risk due to sufficient blinding of experimental information for both participants and experimenters. Similarly, 23 studies were rated as having a low risk of detection bias, as outcome assessors were sufficiently blinded. For attrition bias, 80 studies were assessed as having a low risk because only a small number of samples were lost, or a valid justification for sample loss was provided. Reporting bias was rated as unclear in 102 studies due to the absence of pre-registered study protocols. Other potential sources of bias were also rated as unclear in 104 studies, as information on early study termination, determination of the target sample size, and/or funding sources was insufficient.

**Figure 4.**
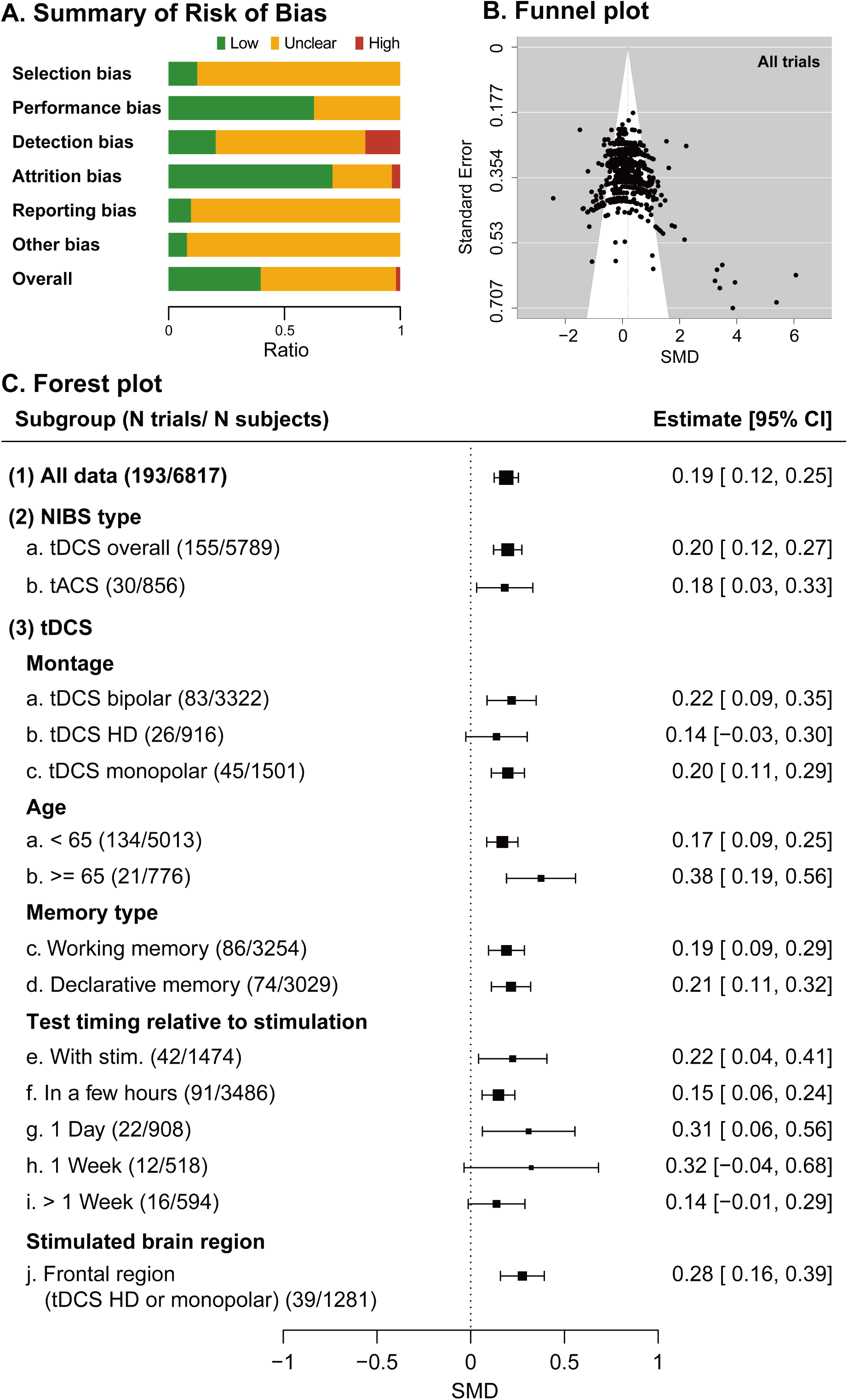
Meta-analyses on the efficacy of NIBS in memory function. (A) Risk of bias assessment. Green, low risk of bias; red, high risk of bias; yellow, unclear. (B) Funnel plot for all data. Each circle represents an individual effect-size estimate plotted against its standard error. (C) Forest plot. Squares show mean effect sizes for each subgroup, with horizontal lines indicating the 95% CI.

### 3.4. Efficacy of NIBS on memory function

A total of 193 trials comparing active stimulation with sham, drawn from 108 articles (total sample size: 6,817 participants), were included in the meta-analysis, which used a multilevel random-effects model. The findings revealed a significant effect of NIBS on memory performance (SMD [95% confidence interval (CI)] = 0.19 [0.12–0.25]; Fig. 4C (1)). However, significant heterogeneity (Q(737) = 1725.45, *p* < .001) and funnel-plot asymmetry were found (Fig. 4B; Begg’s test: τ = 0.09, *p* < .001; Egger’s test: Z = 13.81, *p* < .001).

Further analysis assessed the efficacy of different types of NIBS (Fig. 4C). A significant effect was observed for overall tDCS (SMD [95% CI] = 0.20 [0.12–0.27]) and overall tACS (SMD [95% CI] = 0.18 [0.03–0.33]), with significant heterogeneity (ps < .001); an omnibus moderator test indicated no significant difference between the two modalities (QM(1) = 0.07, p = .788).

Within tDCS, subgroup analyses by electrode montage revealed a significant effect for monopolar tDCS (SMD [95% CI] = 0.20 [0.11–0.29]) and bipolar tDCS (SMD [95% CI] = 0.22 [0.09–0.35]), whereas HD tDCS did not show a significant effect (SMD [95% CI] = 0.14 [-0.03–0.30]); the three montages did not differ significantly from one another (QM(2) = 0.63, p = .731).

In terms of age groups, significant effects were observed in individuals younger than 65 years (SMD [95% CI] = 0.17 [0.09–0.25]) and those aged 65 and older (SMD [95% CI] = 0.38 [0.19–0.56]), and an omnibus moderator test confirmed that the effect was significantly larger in the older group (QM(1) = 17.75, p < .0001). The distribution of effect sizes by age group is shown in Supplementary Figure 2.

In terms of memory type, both working memory (SMD [95% CI] = 0.19 [0.09–0.29]) and declarative memory (SMD [95% CI] = 0.21 [0.11–0.32]) showed significant effects, and the two did not differ significantly (QM(1) = 0.04, p = .842).

We also examined effects according to the timing of memory performance testing relative to tDCS administration. Significant effects were found during stimulation (SMD [95% CI] = 0.22 [0.04–0.41]), within a few hours after stimulation (SMD [95% CI] = 0.15 [0.06–0.24]), and after one day (SMD [95% CI] = 0.31 [0.06–0.56]). By contrast, no significant effects were found at one week (SMD [95% CI] = 0.32 [-0.04–0.68]) or more than one week after stimulation (SMD [95% CI] = 0.14 [-0.01–0.29]). However, an omnibus test across the five timing categories was not significant (QM(4) = 3.13, p = .537), indicating that these differences in significance do not establish a difference between the timing categories themselves.

Regarding stimulation site, to focus on stimulation approaches intended to provide more localised current delivery, we excluded studies using bipolar stimulation and restricted the analysis to HD-tDCS and monopolar tDCS. When these stimulation approaches targeted the frontal region, a significant effect was observed (SMD [95% CI] = 0.28 [0.16–0.39]). Significant heterogeneity was observed across all subgroups, and funnel-plot asymmetry was observed in most cases (see Supplementary Table 2).

### 3.5 Relationship between electric field distribution and memory performance

To identify cortical regions in which tDCS-induced electric fields were associated with greater memory enhancement, we conducted PEC analyses separately for working and declarative memory. The analyses included 65 aggregated effect sizes from 44 WM studies and 60 from 39 DM studies, corresponding to 57 and 51 distinct study-montage pairs, respectively. The mean electric field distributions for both memory types are presented in Fig. S3A and B, with the corresponding vertex-wise E-model t-statistics in Fig. S3C and D.

A positive association surviving correction across 360 ROIs was identified exclusively in the WM E × T model. This association encompassed 100 ROIs, including 71 in the right hemisphere and 29 in the left hemisphere (Fig. 5A). Significant parcels included rh.a9-46v, rh.9-46d, rh.p47r, rh.IFSa, rh.OFC, and rh.11l. The strongest statistical association was observed in right a9-46v, where a one-standard-deviation increase in E × T was associated with a 0.1644 increase in Hedges’ g (t = 8.42, corrected p = 0.0108; Fig. 5B). One standard deviation of E × T corresponded to 1,176.44 V·s/m. No ROIs survived correction in the WM E model or either DM model; the smallest corrected p-values were 0.190, 0.385, and 0.696 for WM E, DM E, and DM E × T, respectively.

**Figure 5.**
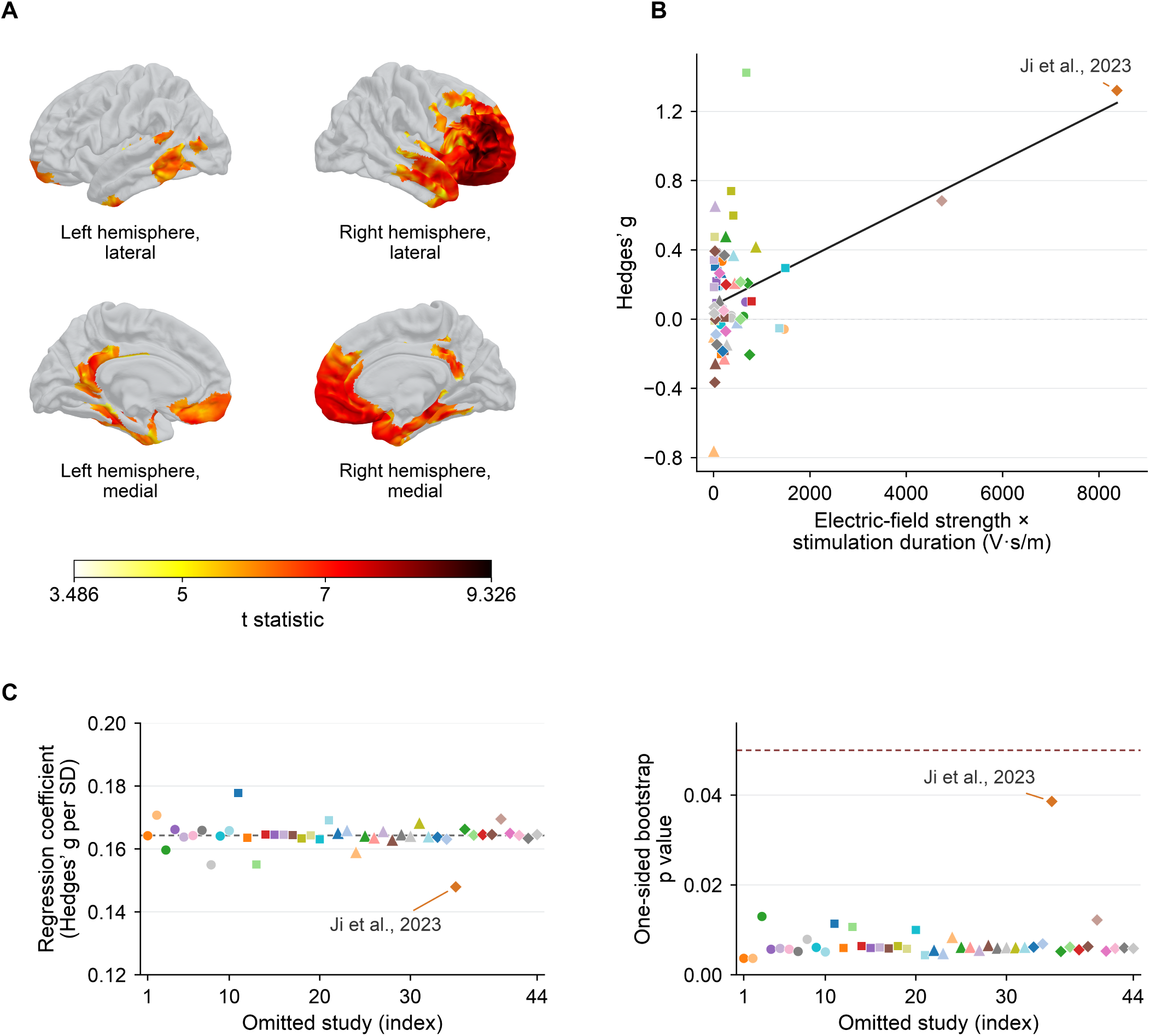
Regional electric-field dose and working-memory effect sizes. (A) Vertex-wise t statistics within ROIs showing significant positive associations between electric-field strength × stimulation duration and Hedges’ g (one-sided p < 0.05, corrected for multiple comparisons across 360 ROIs). The colour scale starts at 3.486, the lowest vertex-wise t statistic within these significant ROIs. (B) Association in the right a9-46v ROI across 65 aggregated observations from 44 working-memory studies. The black line shows adjusted predictions averaged over the observed covariate distribution. Colours and marker shapes identify studies consistently across B and C. (C) Leave-one-study-out analysis of the right a9-46v. Left: regression coefficients per full-sample standard deviation of the predictor, with the dashed line indicating the full-sample estimate. Right: one-sided bootstrap p-values for the fixed ROI, with the dashed line indicating p = 0.05. Each index represents removal of one study and all its observations.

Leave-one-study-out analyses yielded positive coefficients in all 4,400 regressions across the 100 significant ROIs. In right a9-46v, the positive association remained significant in all 44 individual bootstrap tests after study exclusion (β = 0.148–0.178; p = 0.0037–0.0386; Fig. 5C). Exclusion of Ji et al. (2023), which contributed the largest E × T value in this ROI, produced the smallest coefficient and largest individual p-value (β = 0.148, t = 3.39, p = 0.0386). These sensitivity analyses support a robust positive E × T association within WM.

### 3.6. Adverse events

Adverse events related to NIBS were assessed in 32 trials from 22 articles using scale-based indicators, such as the visual analogue scale or numerical rating scale, and 20 trials from 17 articles as the number of cases. Of these, 29 trials from 20 articles (734 active and 460 sham participants) reported means and standard deviations for both groups and entered the analysis of scale-based indicators, and 8 trials from 5 articles (164 active and 109 sham participants) entered the analysis of case numbers, after excluding trials that reported only a global count, did not report counts by symptom, or recorded no events in either group. None of the trials reported serious adverse events.

The analysis of scale-based indicators revealed that the tingling or itching sensation (SMD [95% CI] = 0.20 [0.06–0.34]) was significantly stronger in the active group than in the control group (Fig. 6A). Other measures did not show statistically significant differences.

**Figure 6.**
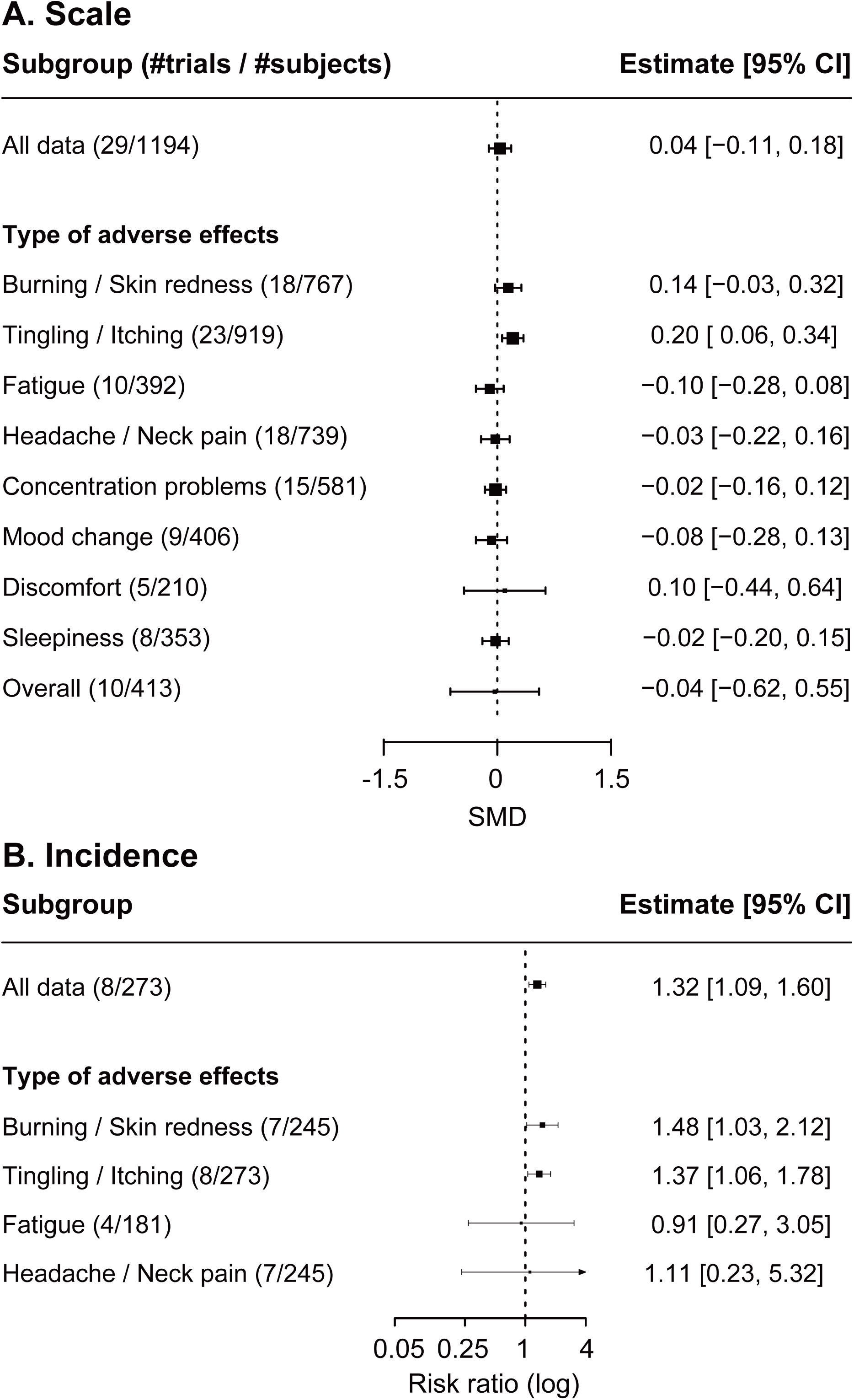
Meta-analyses of adverse events. (A) Forest plot for scale-based indicators. Squares show the mean effect sizes (SMD) for each type of adverse effect, with horizontal lines indicating the 95% CI; positive values indicate more severe symptoms under active stimulation. (B) Forest plot for incidence. Squares show the pooled risk ratio, plotted on a logarithmic scale, and horizontal lines the 95% confidence interval; values greater than 1 indicate more frequent events under active stimulation.

In terms of the number of cases (Fig. 6B), pooling all symptom-specific reports, the incidence was 20% in the active group (95% CI [13–30%]) and 15% in the sham group (95% CI [11–20%]); this is not the proportion of participants with at least one adverse event. Risk ratios were estimated after excluding trials in which no events occurred in either group, and indicated a significantly elevated overall risk under active stimulation (risk ratio [95% CI] = 1.32 [1.09–1.60]). Tingling/itching sensations were the most frequently reported symptom (active: 34% [28–40%]; sham: 24% [19–29%]) and were significantly more likely under active stimulation (risk ratio [95% CI] = 1.37 [1.06–1.78]), as were burning or stimulation-site sensations (active: 23% [9–47%]; sham: 15% [6–33%]; risk ratio [95% CI] = 1.48 [1.03–2.12]). By contrast, fatigue (active: 6% [2–16%]; sham: 5% [2–12%]; risk ratio [95% CI] = 0.91 [0.27–3.05]) and headache or neck pain (active: 11% [6–19%]; sham: 7% [3–16%]; risk ratio [95% CI] = 1.11 [0.23–5.32]) did not differ significantly between groups.

## 4. Discussion

The present review provides a broad assessment of how tES has been used to influence memory performance in healthy adults. Across the included parallel-group design RCT studies, active stimulation produced a small positive pooled effect, equivalent to approximately one-fifth of a standard deviation. Positive estimates were observed for both working and declarative memory, in younger and older adults, and for assessments conducted during stimulation or within approximately 24 h. The subgroup of tDCS targeting the frontal brain region also yielded a positive pooled estimate. These findings indicate that tES can modify performance on laboratory memory tasks, but they do not support uniform enhancement across protocols or individuals. Substantial heterogeneity and funnel-plot asymmetry were present in most analyses, indicating that the results conceal considerable variation and may be influenced by small-study effects or selective publication.

Comparison with previous meta-analyses indicates a broadly consistent pattern of small, context-dependent effects of tES on memory. The present working-memory estimate is comparable to earlier reports of modest benefits from prefrontal tDCS (Brunoni & Vanderhasselt, 2014; Hill et al., 2015; Mancuso et al., 2016; Wischnewski et al., 2021). By restricting the analysis to parallel-group trials, the present review reduces potential carry-over and repeated-testing effects associated with crossover designs, which also explains the smaller evidence base relative to Wischnewski et al. (2021). The positive but imprecise tACS estimate is likewise consistent with Booth et al. (2021), who highlighted substantial protocol-dependent variability.

The evidence was most extensive for tDCS, which produced a comparatively precise estimate because it accounted for most of the included trials. tACS also showed a positive pooled effect, but with a wider confidence interval. The tACS literature combined protocols that differed in frequency, phase relationship, electrode configuration, stimulation timing, and whether stimulation was matched to individual oscillatory activity. Such variation may obscure effects specific to particular oscillatory mechanisms. The non-significant estimate for HD-tDCS may likewise reflect limited evidence, variation among focal montages, or a genuinely smaller effect.

tDCS on the frontal region was the dominant approach and showed a clear positive association with memory performance. The DLPFC was the most frequent nominal target, consistent with its involvement in working-memory maintenance, manipulation, and executive attention (Curtis & D’Esposito, 2003; Kane & Engle, 2002). However, the cortical regions affected by tDCS depend on the complete electrode montage rather than on a single nominal target, and memory-task performance also relies on sustained attention, task-set maintenance, strategic organisation, response monitoring, and resistance to interference. The comparable pooled estimates for working and declarative memory, despite their partly distinct mnemonic mechanisms, are therefore compatible with the possibility that tES influenced domain-general processes shared across tasks rather than selectively strengthening one memory system.

The PEC analysis identified a positive association between electric field strength multiplied by stimulation duration (E × T) and working-memory effect sizes across 100 ROIs, including 71 in the right hemisphere and 29 in the left, with the strongest association in right a9-46v. This analysis retained all age groups and was restricted to assessments during stimulation or within 24 hours. This distribution is more compatible with involvement of distributed frontal regions than with an effect confined to the DLPFC. Engagement of this network could influence broader cognitive performance. Compared with the lower-DLPFC association identified by Wischnewski et al. (2021), the broader distribution observed here may reflect differences in included study designs (we included only parallel-group trials), memory outcomes, montage composition, head models, and spatial inference procedures. The corrected association was identified only in the working-memory E × T model. No ROIs survived correction in the working-memory E model or either declarative-memory model. It should also be noted that PEC is a correlational analysis across studies and montages, and does not demonstrate the causal effects of each region. The concentration of montages over frontal sites and the use of a single template head model also limit anatomical generalisation.

Although similar positive effects were observed across most subgroups (stimulation type, age, memory type, and the test timing relative to stimulation), the current evidence does not establish that tES improves memory in a general sense. Overall, studies are concentrated on a narrow range of laboratory paradigms: most working-memory studies relied on n-back or related tasks, and outcomes were typically assessed using visually presented, low-information stimuli shortly after stimulation. These tasks capture only limited components of memory and also depend on attention, strategy, and response control. Auditory and multisensory memory, associative and episodic learning, naturalistic event memory, and ecologically relevant outcomes remain comparatively underrepresented. It therefore remains unclear whether the observed effects generalise to everyday remembering, academic learning, or untrained cognitive abilities.

Pooled effects were significant in both younger and older adults and numerically larger in older adults. However, fewer studies included older adults, and these estimates alone do not establish greater efficacy in this group. Several factors may contribute to this apparent age-related difference. High baseline memory performance in younger adults may limit measurable improvement through ceiling effects, whereas age-related reductions in performance may leave greater room for improvement in older adults. Studies in older adults also involved more repeated stimulation sessions and longer intervention protocols, suggesting that differences in treatment exposure may partly account for their larger pooled effect. Furthermore, funnel-plot asymmetry was more pronounced in the older-adult subgroup (Supplementary Figure 2), raising greater concern about publication bias in this group. Expectations of preventive benefits could potentially favour the selective publication of positive findings, although the present analyses cannot establish the reasons for this asymmetry. Thus, the larger pooled effect in older adults should be interpreted cautiously rather than taken as evidence that tES is intrinsically more effective in this population.

The durability of these effects also remains uncertain. Effects were most consistently observed during stimulation or within the following day, whereas longer-term estimates were imprecise. Evidence therefore remains limited on whether tES can help maintain memory function during healthy ageing before clinical impairment emerges. Studies with delayed follow-up assessments, larger samples of healthy older adults, and a broader range of memory tasks are needed to determine whether the observed benefits persist and translate into practically meaningful improvements.

The adverse-event findings indicate that tES was generally well tolerated under the studied protocols. No serious adverse events were reported, and the clearest active–sham difference was a modest increase in tingling or itching. However, adverse events were systematically quantified in only a subset of trials, and follow-up was usually short. The absence of reported serious events is not equivalent to comprehensive evidence of safety, particularly for repeated or unsupervised use. Scalp sensations may also compromise blinding and permit expectancy effects; future trials should assess both adverse events and blinding success using standardised procedures.

Taken together, the accumulating evidence suggests that tES can produce a small improvement in memory-task performance, although substantial heterogeneity and small-study effects limit the certainty and generalisability of this estimate. The field should therefore move beyond repeatedly asking whether tES has any effect and focus increasingly on identifying the stimulation, task, and participant conditions under which benefits become reliable, durable, and practically meaningful. This will require direct comparisons of candidate moderators and systematic links between stimulation parameters, cortical electric fields, cognitive processes, and behavioural outcomes.

The evidence base has expanded rapidly and shows some signs of methodological maturation, but important weaknesses remain. Just over half of the included articles (57 of 113) were published between 2022 and 2025, and the year-by-year screening pattern suggested that a greater proportion of recent reports met the present eligibility criteria. Performance and attrition bias were also commonly judged low. However, eligibility is only an indirect indicator of study quality, and risk-of-bias judgements remained frequently unclear because randomisation, allocation concealment, outcome-assessor blinding, and preregistered analysis plans were inadequately reported. Data from many studies also remained unavailable, while incomplete reporting of electrode positions, dimensions, and materials prevented some otherwise eligible studies from being included in the PEC analysis. Future studies should therefore use adequately powered and preregistered designs, specify primary outcomes and analysis plans in advance, report randomisation and blinding procedures transparently, provide sufficient information to reconstruct the complete stimulation montage, and share data and analysis code. Such improvements are essential for distinguishing reproducible moderators of efficacy from variability arising from methodological and reporting differences.

## 5. Conclusion

tES was associated with small improvements in working- and declarative-memory task performance. Positive pooled effects were observed during stimulation and within approximately 24 hours; longer-term effects remain uncertain. However, to achieve a more generalisable understanding of NIBS’s effectiveness in memory enhancement, addressing biases in research themes is crucial. Many existing studies disproportionately focus on specific memory types, sensory modalities, memory tasks, and adverse effects, leaving significant gaps in knowledge. Future research should broaden its scope by investigating underexplored areas, such as long-term memory, non-visual sensory modalities for stimulus presentation, and alternative electrode montages. A more balanced and comprehensive exploration of these variables will contribute to a deeper understanding of how NIBS modulates memory across diverse conditions.

## Authorship contribution statement

**Yujin Goto:** Methodology, Software, Formal analysis, Data Curation, Visualisation, Writing **–** original draft, Writing – review & editing.

**Ryoji Onagawa:** Methodology, Software, Formal analysis, Data Curation, Visualisation, Writing **–** original draft, Writing – review & editing.

**Mitsuaki Takemi:** Formal analysis, Visualisation, Writing **–** original draft, Funding acquisition, Writing – review & editing.

**Keiichi Ishikawa:** Data Curation, Validation, Writing – review & editing.

**Suzuka Narukawa:** Data Curation, Validation, Writing – review & editing.

**Kaoru Amano:** Conceptualisation, Validation, Writing – review & editing, Supervision.

**Koichi Hosomi:** Conceptualisation, Methodology, Validation, Writing – review & editing, Supervision, Project administration.

## Data and code availability statement

All data used in this review were obtained from original publications and shared manuscripts by the authors. The full version of the metadata upon which the meta-analyses were based and the code are available in the OSF repository (https://osf.io/rfu3n/).

## Acknowledgments

This work was conducted as a part of Braintech Guidebook development in JST Moonshot R&D (Grant Number JPMJMS2012) to Mitsuaki Takemi. We thank the members of the Evidence Evaluation Committee for Braintech Guidebook, a specially organized group for the Moonshot R&D project, for their comments on the manuscript.

## Declaration of generative AI and AI-assisted technologies in the manuscript preparation process

During the preparation of this manuscript, the authors used ChatGPT (OpenAI) and Claude (Anthropic) to assist with data checking and English language editing. The authors reviewed and verified the AI-assisted outputs and take full responsibility for the content of the manuscript.

