## Supplementary File 1 for "Transcranial electrical stimulation for memory enhancement: A systematic review and meta-analysis"

Supplementary File 1. Characteristics of the included studies. Participant counts, memory outcomes, and stimulation parameters for each study and group are shown.

| Study and participants | | | | | | Memory outcomes | | | | Brain stimulation | | | | | | |
|---|---|---|---|---|---|---|---|---|---|---|---|---|---|---|---|---|
| Reference | Total N (female/male) | Group | Group N | Female | Male | Age group | Memory type | Recall or recognition | Outcome | NIBS type | Montage | Anode (tDCS) or electrode 1 | Return or other electrode(s) | Duration per session (min) | Total sessions | Amplitude (as reported) |
| (Aktürk et al., 2022) | 46 (35/11) | ITF | 15 | 13 | 2 | Adult | Declarative, Working | Recall | Accuracy | tACS | BP-tACS | F3 | P3 | 20 | 1 | 1.5 (peak-to-peak) |
| | | ITF-1 | 16 | 12 | 4 | | | | | tACS | BP-tACS | F3 | P3 | 20 | 1 | 1.5 (peak-to-peak) |
| | | Control | 15 | 10 | 5 | | | | | Sham | | F3 | P3 | | 1 | |
| (Biel et al., 2022) | 46 (28/18) | Anti-phase | 11 | NR | NR | Adult | Working | Recognition | Accuracy, RT | tACS | BP-tACS | F3 | P3 | 14 | 1 | 2 |
| | | In-phase Cz | 11 | NR | NR | | | | | tACS | BP-tACS | F3 and P3 | Cz | 14 | 1 | 2 |
| | | In-phase focal | 12 | NR | NR | | | | | tACS | BP-tACS | F3 and P3 | F7, Fz, C3, P7, Pz | 14 | 1 | 3 |
| | | Control | 12 | NR | NR | | | | | Sham | | F3 | P3 | | 1 | |
| (Luckey et al., 2022) | 85 (49/36) | tDCS | 25 | | | Adult | Declarative | Recall | Accuracy | tDCS | BP-tDCS | left C2 nerve dermatome (left occipital region) | right C2 nerve dermatome (right occipital region) | 4.16667 | 3 | 1.5 |
| | | 40Hz tACS | 24 | | | | | | | tACS | BP-tACS | left C2 nerve dermatome (left occipital region) | right C2 nerve dermatome (right occipital region) | 4.16667 | 3 | 3 (peak to peak) |
| | | Control (1Hz tACS) | 11 | | | | | | | tACS | BP-tACS | left C2 nerve dermatome (left occipital region) | right C2 nerve dermatome (right occipital region) | 4.16667 | 3 | 3 |
| | | Control (sham) | 25 | | | | | | | Sham | | left C2 nerve dermatome (left occipital region) | right C2 nerve dermatome (right occipital region) | | 3 | |
| (Antonenko et al., 2022) | 51 (36/15) | Intervention | 24 | | | Aged | Working, Declarative, Others | Recall, Recognition | Accuracy | tDCS | BP-tDCS | F3 | FP2 | 20 | 1 | 1 |
| | | Control | 27 | | | | | | | Sham | | F3 | FP2 | | 1 | |
| (Teixeira-Santos et al., 2022) | 54 (-/-) | Intervention | 18 | NR | NR | Aged | Working | Recall | Accuracy | tDCS | BP-tDCS | F3 | FP2 | 20 | 5 | 2 |
| | | Control | 18 | NR | NR | | | | | Sham | BP-tDCS | F3 | FP2 | | 5 | |
| | | Control | 18 | NR | NR | | | | | Sham | | F3 | FP2 | | 5 | |
| (Brambilla et al., 2021) | 47 (21/26) | Intervention 1 mA | 14 | 6 | 8 | Aged | Declarative, Working | Recall, Recognition | Accuracy | tRNS | BP-tRNS | F3 | F4 | 20 | 5 | 1 |
| | | Intervention 0.705 mA | 14 | 7 | 7 | | | | | tRNS | BP-tRNS | F3 | F4 | 20 | 5 | 0.705 |
| | | Control | 19 | 8 | 11 | | | | | Sham | | F3 | F4 | | 5 | |
| (Meléndez et al., 2021) | 29 (18/11) | Intervention | 16 | 10 | 6 | Aged | Declarative | Recognition | Accuracy | tDCS | BP-tDCS | F7 | FP2 | 20 | 1 | 2 |
| | | Control | 13 | 8 | 5 | | | | | Sham | | F7 | FP2 | | 1 | |
| (Fernández et al., 2021) | 23 (-/-) | Intervention | 12 | NR | NR | Adult | Declarative, Working | Recall, Recognition | Accuracy, RT | tDCS | BP-tDCS | F3 | FP2 | 18 | 2 | 2 |
| | | Control | 11 | NR | NR | | | | | Sham | | F3 | FP2 | | 2 | |
| (Gilson et al., 2021) | 69 (41/28) | left tDCS | 16 | 10 | 6 | Adult | Declarative | Recall | Accuracy | tDCS | BP-tDCS | F3 | F4 | 20 | 1 | 1 |
| | | right tDCS | 17 | 11 | 6 | | | | | tDCS | BP-tDCS | F4 | F3 | 20 | 1 | 1 |
| | | Control | 20 | 11 | 9 | | | | | Sham | BP-tDCS | F3 | F4 | | 1 | |
| | | Control (TMR-only) | 16 | 9 | 7 | | | | | No stimulation | | | | | | |
| (Dubravac et al., 2020) | 60 (34/26) | Intervention, left | 20 | NR | NR | Adult | Declarative | Recognition | Accuracy | tDCS | BP-tDCS | P3 | P6 | 20 | 1 | 1 |
| | | Intervention, right | 20 | NR | NR | | | | | tDCS | BP-tDCS | P6 | P3 | 20 | 1 | 1 |
| | | Control | 20 | NR | NR | | | | | Sham | | P3 | P6 | | 1 | |
| (Alonso et al., 2021) | 75 (64/11) | Anodal | 25 | 20 | 5 | Adult | Declarative | Recognition | Accuracy | tDCS | MP-tDCS | FT10 | left shoulder | 20 | 1 | 2 |
| | | Cathodal | 25 | 22 | 3 | | | | | tDCS | MP-tDCS | left shoulder | FT10 | 20 | 1 | 2 |
| | | Control | 25 | 22 | 3 | | | | | Sham | | FT10 | left shoulder | | 1 | |
| (Byrne et al., 2020) | 48 (31/17) | BDR training + Intervention tDCS | 16 | 9 | 7 | Adult | Working | Recall, Recognition | Accuracy | tDCS | BP-tDCS | F3 | right supraorbital | 10 | 3 | 1 |
| | | BDR training + Sham | 16 | 11 | 5 | | | | | Sham | BP-tDCS | F3 | right supraorbital | | 3 | |
| | | Visual search training + Sham | 16 | 11 | 5 | | | | | Sham | | F3 | right supraorbital | | 3 | |
| (Firouzkouhi Moghadam et al., 2020) | 45 (22/23) | left DLPFC | 15 | NR | NR | Adult | Working, Declarative | Recall, Recognition | Accuracy, RT | tDCS | BP-tDCS | F3 | right supraorbital | | 5 | 2 |
| | | right DLPFC | 15 | NR | NR | | | | | tDCS | BP-tDCS | F4 | left supraorbital | | 5 | 2 |
| | | Control | 15 | NR | NR | | | | | Sham | | F3 | right supraorbital | | 5 | |

| Study and participants | | | | | | Memory outcomes | | | | Brain stimulation | | | | | | |
|---|---|---|---|---|---|---|---|---|---|---|---|---|---|---|---|---|
| Reference | Total N (female/male) | Group | Group N | Female | Male | Age group | Memory type | Recall or recognition | Outcome | NIBS type | Montage | Anode (tDCS) or electrode 1 | Return or other electrode(s) | Duration per session (min) | Total sessions | Amplitude (as reported) |
| (Petrovskaya et al., 2020) | 97 (-/-) | DLPFC online | 24 | NR | NR | Adult | Declarative | Recognition | Accuracy | tDCS | MP-tDCS | F3 | left shoulder | 10 | 1 | 2 |
| | | DLPFC offline | 25 | NR | | | | | | tDCS | MP-tDCS | F3 | left shoulder | 10 | 1 | 2 |
| | | VLPFC online | 23 | NR | | | | | | tDCS | MP-tDCS | F7 | left shoulder | 10 | 1 | 2 |
| | | Control | 25 | NR | | | | | | Sham | | F3 | left shoulder | | 1 | |
| (Jones et al., 2020) | 23(11/12) | Intervention | 11 | 5 | 6 | Adult | Working | Recognition | Accuracy | tDCS | MP-tDCS | F4 or P4 | left cheek | 15 | 4 | 1.5 |
| | | Control | 12 | 6 | 6 | | | | | Sham | | F4 or P4 | left cheek | | 4 | |
| (Zhu et al., 2020) | 51 (31/20) | PPC | 17 | 10 | 7 | Adult | Working | Recognition | Accuracy | tDCS | MP-tDCS | P3 | right cheek | 20 | 1 | 1.5 |
| | | IFG | 17 | 10 | 7 | | | | | tDCS | MP-tDCS | crossing point between F7-Cz and T3-Fz | right cheek | 20 | 1 | 1.5 |
| | | Control | 17 | 11 | 6 | | | | | Sham | | P3 or crossing point between F7-Cz and T3-Fz (half each) | right cheek | | 1 | |
| (Perceval et al., 2020) | 101 (75/26) | Adult intervention | 21 | 13 | 8 | Adult | Working, Declarative, Others | Recall, Recognition | Accuracy | tDCS | BP-tDCS | (a) the intersection of T3-F3 and F7-C3 and (b) the midpoint between F7-F3. -> center of a line connecting points (a) and (b). | right supraorbital | 20 | 5 | 1 |
| | | Older intervention | 30 | 26 | 4 | Aged | | | | tDCS | BP-tDCS | (a) the intersection of T3-F3 and F7-C3 and (b) the midpoint between F7-F3. -> center of a line connecting points (a) and (b). | right supraorbital | 20 | 5 | 1 |
| | | Adult control | 20 | 12 | 8 | Adult | | | | Sham | BP-tDCS | (a) the intersection of T3-F3 and F7-C3 and (b) the midpoint between F7-F3. -> center of a line connecting points (a) and (b). | right supraorbital | | 5 | |
| | | Older control | 30 | 24 | 6 | Aged | | | | Sham | | (a) the intersection of T3-F3 and F7-C3 and (b) the midpoint between F7-F3. -> center of a line connecting points (a) and (b). | right supraorbital | | 5 | |
| (Huo et al., 2020) | 49 (20/29) | Intervention | 24 | 11 | 13 | Aged | Declarative | Recognition | Accuracy, Others | tDCS | MP-tDCS | F3 | right deltoid | 30 | 10 | 2 |
| | | Control | 25 | 9 | 16 | | | | | Sham | | F3 | right deltoid | | 10 | |
| (Mainz et al., 2020) | 75 (0/75) | Anodal | 25 | 0 | 25 | Adult | Declarative | Recall | Accuracy | tDCS | BP-tDCS | FPz | between Inion and Oz | 20 | 1 | 2 |
| | | Cathodal | 25 | 0 | 25 | | | | | tDCS | BP-tDCS | between Inion and Oz | FPz | 20 | 1 | 2 |
| | | Control | 25 | 0 | 25 | | | | | Sham | | FPz | between Inion and Oz | | 1 | |
| (Baumert et al., 2020) | 87 (-/-) | Anodal | 26 | NR | NR | Adult | Working | Recognition | Accuracy, RT | tDCS | MP-tDCS | F3 | right upper trapezius | 20 | 1 | 1 |
| | | Cathodal | 31 | NR | NR | | | | | tDCS | MP-tDCS | right upper trapezius | F3 | 20 | 1 | 1 |
| | | Control | 30 | NR | NR | | | | | Sham | | F3 | right upper trapezius | | 1 | |

| Study and participants | | | | | | Memory outcomes | | | | Brain stimulation | | | | | | |
|---|---|---|---|---|---|---|---|---|---|---|---|---|---|---|---|---|
| Reference | Total N (female/male) | Group | Group N | Female | Male | Age group | Memory type | Recall or recognition | Outcome | NIBS type | Montage | Anode (tDCS) or electrode 1 | Return or other electrode(s) | Duration per session (min) | Total sessions | Amplitude (as reported) |
| (H. Lu et al., 2021) | 43 (19/24) | Intervention | 22 | 10 | 12 | Adult | Working | Recognition | Accuracy | tDCS | HD-tDCS | F3 | AF3, F1, F5, FC3 | 20 | 9 | 1.5 |
| | | Control | 21 | 9 | 12 | | | | | Sham | | F3 | AF3, F1, F5, FC3 | | 9 | |
| (Marrero et al., 2020) | 65 (56/9) | Anodal | 24 | NR | NR | Adult | Declarative | Recognition | Accuracy | tDCS | MP-tDCS | T8 | left shoulder | 20 | 1 | 2 |
| | | Cathodal | 21 | NR | NR | | | | | tDCS | MP-tDCS | left shoulder | T8 | 20 | 1 | 2 |
| | | Control | 20 | NR | NR | | | | | Sham | | T8 | left shoulder | | 1 | |
| (Berlin et al., 2020) | 44 (31/13) | Intervention | 22 | 16 | 6 | Adult | Working | Recognition | Accuracy | tDCS | MP-tDCS | F3 | left upper trapezius | 30 | 1 | 1 |
| | | Control | 22 | 15 | 7 | | | | | Sham | | F3 | left upper trapezius | | 1 | |
| (Bashir et al., 2019) | 36 (16/20) | Intervention | 18 | NR | NR | Adult | Declarative | Recognition | Accuracy | tDCS | BP-tDCS | F3 | C4 | 20 | 1 | 2 |
| | | Control | 18 | NR | NR | | | | | Sham | | F3 | C4 | | 1 | |
| (Medvedeva et al., 2019) | 120 (-/-) | Online during encoding | 17 | NR | NR | Adult | Declarative | Recognition | Accuracy | tDCS | MP-tDCS | F7 | right deltoid | 9 | 1 | 2 |
| | | Offline | 15 | NR | NR | Adult | | | | tDCS | MP-tDCS | F7 | right deltoid | 10 | 1 | 2 |
| | | Online during retrieval | 16 | NR | NR | Adult | | | | tDCS | MP-tDCS | F7 | right deltoid | 10 | 1 | 2 |
| | | Offline | 15 | NR | NR | Adult | | | | tDCS | MP-tDCS | F7 | right deltoid | 10 | 1 | 2 |
| | | Online during encoding | 11 | NR | NR | Aged | | | | tDCS | MP-tDCS | F7 | right deltoid | 9 | 1 | 2 |
| | | Control during encoding | 17 | NR | NR | Adult | | | | Sham | MP-tDCS | F7 | right deltoid | | 1 | |
| | | Control during retrieval | 18 | NR | NR | Adult | | | | Sham | MP-tDCS | F7 | right deltoid | | 1 | |
| | | Control during encoding | 11 | NR | NR | Aged | | | | Sham | | F7 | right deltoid | | 1 | |
| (Wang et al., 2019) | 20 (11/9) | Intervention | 19 | NR | NR | Adult | Working | Recognition | Others | tDCS | HD-tDCS | F3 | Fz, FP1, FT7, C3 | 30 | 10 | 2 |
| | | Control | 19 | NR | NR | | | | | Sham | | F3 | Fz, FP1, FT7, C3 | | 10 | |
| (Mashal & Metzuyanim-Gorelick, 2019) | 25 (11/14) | Intervention | 16 | 8 | 8 | Adult | Working | Recognition | Accuracy, RT | tDCS | BP-tDCS | F3 | F4 | 20 | 6 | 2 |
| | | Control | 9 | 3 | 6 | | | | | Sham | | F3 | F4 | | 6 | |
| (Lang et al., 2019) | 59 (31/28) | tDCS | 21 | 12 | 9 | Adult | Declarative | Recognition | Accuracy | tDCS | HD-tDCS | P10 | FP1,P2,P3,PO7 | 10 | 1 | 2 |
| | | tACS | 19 | 7 | 12 | | | | | tACS | HD-tACS | P10 | FP1,P2,P3,PO7 | 10 | 1 | 4 |
| | | Control | 19 | 12 | 7 | | | | | Sham | | P10 | FP1,P2,P3,PO7 | | 1 | |
| (Naka et al., 2018) | 20 (10/10) | Intervention | 10 | 5 | 5 | Adult | Working | Recognition | Accuracy, RT | tDCS | HD-tDCS | F3 | C3,F7,FP1,Fz | 16 | 1 | 1.5 |
| | | Control | 10 | 5 | 5 | | | | | Sham | | F3 | C3,F7,FP1,Fz | | 1 | |
| (Nikolin et al., 2019) | 78 (51/27) | DLPFC | 26 | 18 | 8 | Adult | Working | Recognition | Accuracy, RT | tDCS | HD-tDCS | F3 | F5,AF3,F1,FC3 | 20 | 1 | 2 |
| | | IPS | 26 | 17 | 9 | | | | | tDCS | HD-tDCS | P3 | P7,Pz,C3,O1 | 20 | 1 | 2 |
| | | Control | 26 | 16 | 10 | | | | | Sham | | F3 or P3 (randomized) | F5, AF3, F1, FC3 or P7, Pz, C3, O1 (corresponding montage) | | 1 | |
| (Nikolin et al., 2018) | 99 (52/47) | 1 mA tDCS | 20 | 12 | 8 | Adult | Working | Recognition | Accuracy, RT | tDCS | BP-tDCS | F3 | F4 | 15 | 1 | 1 |
| | | 2 mA tDCS | 20 | 12 | 8 | | | | | tDCS | BP-tDCS | F3 | F4 | 15 | 1 | 2 |
| | | Sham1 (0.016 mA background; 2 mA ramp) | 20 | 8 | 12 | | | | | Sham | | F3 | F4 | | 1 | |
| | | Sham2 (0.034 mA background; 1 mA ramp) | 20 | 9 | 11 | | | | | Sham | | F3 | F4 | | 1 | |

| Study and participants | | | | | | Memory outcomes | | | | Brain stimulation | | | | | | |
|---|---|---|---|---|---|---|---|---|---|---|---|---|---|---|---|---|
| Reference | Total N (female/male) | Group | Group N | Female | Male | Age group | Memory type | Recall or recognition | Outcome | NIBS type | Montage | Anode (tDCS) or electrode 1 | Return or other electrode(s) | Duration per session (min) | Total sessions | Amplitude (as reported) |
| | | Off (electrode leads unplugged) | 19 | 11 | 8 | | | | | Off-stimulation mode | | F3 | F4 | 0 | 1 | 0 |
| (Arciniega et al., 2018) | 36 (23/13) | Intervention | NR | NR | NR | Adult | Working | Recognition | Accuracy | tDCS | MP-tDCS | P4 | left cheek | 10 | 1 | 1.5 |
| | | Control | NR | NR | NR | | | | | Sham | | P4 | left cheek | | 1 | |
| (Brunyé et al., 2018) | 50 (17/33) | Intervention | 25 | 8 | 17 | Adult | Declarative | Recognition | Accuracy, RT | tDCS | Other-tDCS | Fp1, Fp2 | AF3, F4, P8 | 20 | 1 | 1.5 |
| | | Control | 25 | 9 | 16 | | | | | Sham | Other-tDCS | Fp1, Fp2 | AF3, F4, P8 | | 1 | |
| (Díez et al., 2017) | 65 (48/17) | Anodal | 22 | NR | NR | Adult | Declarative | Recognition | Accuracy | tDCS | MP-tDCS | FT9 | right shoulder | 20 | 1 | 2 |
| | | Cathodal | 21 | NR | NR | | | | | tDCS | MP-tDCS | right shoulder | FT9 | 20 | 1 | 2 |
| | | Control | 22 | NR | NR | | | | | Sham | | FT9 | right shoulder | | 1 | |
| (Gaynor & Chua, 2016) | 72 (41/31) | PFC | 24 | | | Adult | Declarative | Recognition | Accuracy | tDCS | BP-tDCS | F3 | right supraorbital | 20 | 1 | 2 |
| | | Parietal | 24 | | | | | | | tDCS | BP-tDCS | CP3 | CP4 | 20 | 1 | 2 |
| | | Control | 24 | | | | | | | Sham | | | | | 1 | |
| (Leshikar et al., 2017) | 42 (26/16) | Intervention | 21 | 15 | 6 | Adult | Declarative | Recall, Recognition | Accuracy, Others | tDCS | MP-tDCS | F3 | right deltoid | 25 | 1 | 1.5 |
| | | Control | 21 | 11 | 10 | | | | | Sham | | F3 | right deltoid | | 1 | |
| (Jones et al., 2017) | 23 (-/-) | Intervention | 11 | NR | NR | Adult | Working | Recognition | Accuracy | tDCS | MP-tDCS | F4 or P4 | left cheek | 15 | 4 | 1.5 |
| | | Control | 12 | 6 | 6 | | | | | Sham | | F4 or P4 | left cheek | | 4 | |
| (Trumbo et al., 2016) | 72 (36/36) | spatial WM left DLPFC | 12 | 6 | 6 | Adult | Working | Recognition | Accuracy, RT | tDCS | MP-tDCS | F3 | area near the bicep between the elbow and shoulder (contralateral to the stimulating hemisphere) | 30 | 1 | 2 |
| | | spatial WM right DLPFC | 12 | 6 | 6 | | | | | tDCS | MP-tDCS | F4 | area near the bicep between the elbow and shoulder | 30 | 1 | 2 |
| | | verbal WM left DLPFC | 12 | 6 | 6 | | | | | tDCS | MP-tDCS | F3 | area near the bicep between the elbow and shoulder | 30 | 1 | 2 |
| | | verbal WM right DLPFC | 12 | 6 | 6 | | | | | tDCS | MP-tDCS | F4 | area near the bicep between the elbow and shoulder | 30 | 1 | 2 |
| | | spatial WM control | 12 | 6 | 6 | | | | | Sham | | F3 or F4 | area near the bicep between the elbow and shoulder, contralateral to the sham electrode | | 1 | |
| | | verbal WM Cotrol | 12 | 6 | 6 | | | | | Sham | | F3 or F4 | area near the bicep between the elbow and shoulder, contralateral to the sham electrode | | 1 | |
| (Stephens & Berryhill, 2016) | 90 (49/41) | Intervention 1mA | 30 | 16 | 14 | Aged | Working | Recall, Recognition | Accuracy | tDCS | MP-tDCS | F4 | left cheek | 15 | 5 | 1 |
| | | Intervention 2mA | 30 | 17 | 13 | | | | | tDCS | MP-tDCS | F4 | left cheek | 15 | 5 | 2 |
| | | Control | 30 | 16 | 14 | | | | | Sham | | F4 | left cheek | | 5 | |
| (Bogdanov & Schwabe, 2016) | 120 (60/60) | Stress condition, anodal | 20 | 10 | 10 | Adult | Working | Recall | Accuracy | tDCS | BP-tDCS | F4 | Cz | 8 | 1 | 1.075 |
| | | Stress condition, cathodal | 20 | 10 | 10 | | | | | tDCS | BP-tDCS | Cz | F4 | 8 | 1 | 1.075 |
| | | Control condition, anodal | 20 | 10 | 10 | | | | | tDCS | BP-tDCS | F4 | Cz | 8 | 1 | 1.075 |

| Study and participants | | | | | | Memory outcomes | | | | Brain stimulation | | | | | | |
|---|---|---|---|---|---|---|---|---|---|---|---|---|---|---|---|---|
| Reference | Total N (female/male) | Group | Group N | Female | Male | Age group | Memory type | Recall or recognition | Outcome | NIBS type | Montage | Anode (tDCS) or electrode 1 | Return or other electrode(s) | Duration per session (min) | Total sessions | Amplitude (as reported) |
| | | Control condition, cathodal | 20 | 10 | 10 | | | | | tDCS | BP-tDCS | Cz | F4 | 8 | 1 | 1.075 |
| | | Stress condition, control | 20 | 10 | 10 | | | | | Sham | | F4 | Cz | | 1 | |
| | | Cotrol condition. control | 20 | 10 | 10 | | | | | Sham | | F4 | Cz | | 1 | |
| (Schaal, Krause, et al., 2015) | 72 (47/25) | Nonmusician cathodal lSMG | 12 | 8 | 4 | Adult | Others | Recall, Recognition | Accuracy | tDCS | BP-tDCS | right supraorbital | CP3 | 20 | 1 | 2 |
| | | Nonmusician cathodal rSMG | 12 | 9 | 3 | | | | | tDCS | BP-tDCS | left supraorbital | CP4 | 20 | 1 | 2 |
| | | Musician cathodal lSMG | 12 | 7 | 5 | | | | | tDCS | BP-tDCS | right supraorbital | CP3 | 20 | 1 | 2 |
| | | Musician cathodal rSMG | 12 | 7 | 5 | | | | | tDCS | BP-tDCS | left supraorbital | CP4 | 20 | 1 | 2 |
| | | Nonmusician control | 12 | 7 | 5 | | | | | Sham | | CP3 | right supraorbital | | 1 | |
| | | Musicians control lSMG | 12 | 9 | 3 | | | | | Sham | | CP3 | right supraorbital | | 1 | |
| (Schaal, Javadi, et al., 2015) | 50 (31/19) | Exp1 anodal right PPC | 13 | 7 | 6 | Adult | Others | Recognition | Accuracy | tDCS | BP-tDCS | P4 | left supraorbital | 15 | 1 | 2 |
| | | Exp2 anodal right PPC | 12 | 8 | 4 | | | | | tDCS | BP-tDCS | P4 | left supraorbital | 15 | 1 | 2 |
| | | Exp2 anodal left PPC | 12 | 8 | 4 | | | | | tDCS | BP-tDCS | P3 | right supraorbital | 15 | 1 | 2 |
| | | Exp1 Sham right PPC | 13 | 8 | 5 | | | | | Sham | | P4 | left supraorbital | | 1 | |
| (Jones et al., 2015) | 72 (49/23) | PFC | 18 | NR | NR | Aged | Working | Recall, Recognition | Accuracy | tDCS | MP-tDCS | F4 | left cheek | 10 | 10 | 1.5 |
| | | PPC | 18 | NR | NR | | | | | tDCS | MP-tDCS | P4 | left cheek | 10 | 10 | 1.5 |
| | | PFC/PPC alternating | 18 | NR | NR | | | | | tDCS | MP-tDCS | F4/P4 | left cheek | 10 | 10 | 1.5 |
| | | Control | 18 | NR | NR | | | | | Sham | | F4 or P4 | left cheek | | 10 | |
| (Park et al., 2014) | 40 (27/13) | Intervention | 20 | 13 | 7 | Aged | Working | Recall, Recognition | Accuracy, RT | tDCS | MP-tDCS | F3, F4 | nondominant arm | 30 | 10 | 2 |
| | | Control | 20 | 14 | 6 | | | | | Sham | | F3, F4 | nondominant arm | | 10 | |
| (Meiron & Lavidor, 2014) | 24 (24/0) | Intervention | 12 | 12 | 0 | Adult | Working | Recognition | RT | tACS | BP-tACS | midpoint of F3 and AF3 | midpoint of F4 and AF4 | 20 | 1 | 1 |
| | | Control | 12 | 12 | 0 | | | | | Sham | | midpoint of F3 and AF3 | midpoint of F4 and AF4 | | 1 | |
| (Richmond et al., 2014) | 58 (39/19) | Intervention + WM training | 20 | 13 | 7 | Adult | Working | Recall | Accuracy | tDCS | BP-tDCS | F3 | F4 | 15 | 10 | 1.5 |
| | | Sham + WM training | 20 | 13 | 7 | | | | | Sham | BP-tDCS | F3 | F4 | | 10 | |
| | | No-contact control | 18 | 13 | 5 | | | | | No stimulation | | | | 0 | 0 | 0 |
| (Meinzer et al., 2013) | 40 (24/16) | Intervention | 20 | NR | NR | Adult | Declarative | Recall, Recognition | Accuracy | tDCS | BP-tDCS | CP5 | right supraorbital | 20 | 5 | 1 |
| | | Control | 20 | NR | NR | | | | | Sham | | CP5 | right supraorbital | | 5 | |
| (Schaal et al., 2013) | 24 (16/8) | Intervention | 12 | 8 | 4 | Adult | Others | Recall, Recognition | Accuracy | tDCS | BP-tDCS | CP3 | right supraorbital | 35-40 | 1 | 2 |
| | | Control | 12 | 8 | 4 | | | | | Sham | | CP3 | right supraorbital | | 1 | |
| (Lally et al., 2013) | 21 (14/7) | Intervention | 10 | NR | NR | Adult | Working | Recognition | Accuracy, RT | tDCS | MP-tDCS | F3 | right cheek | 10 | 2 | 1 |
| | | Control | 11 | NR | NR | | | | | Sham | | F3 | right cheek | | 2 | |
| (Shtoots et al., 2024) | 72 (39/33) | Frontal Theta | 18 | 11 | 7 | Adult | Declarative | Recall | Accuracy | tACS | HD-tACS | Fz | AF4, AF3, FC2, FC1 | 20 | 1 | 1.5 |
| | | Frontal Beta | 18 | 10 | 8 | | | | | tACS | HD-tACS | Fz | AF4, AF3, FC2, FC1 | 20 | 1 | 1.5 |
| | | Parietal Theta | 18 | 9 | 9 | | | | | tACS | BP-tACS | P1–P4 | chin | 20 | 1 | 1.5 |
| | | Control | 18 | 9 | 9 | | | | | Sham | | | | | 1 | |

| Study and participants | | | | | | Memory outcomes | | | | Brain stimulation | | | | | | |
|---|---|---|---|---|---|---|---|---|---|---|---|---|---|---|---|---|
| Reference | Total N (female/male) | Group | Group N | Female | Male | Age group | Memory type | Recall or recognition | Outcome | NIBS type | Montage | Anode (tDCS) or electrode 1 | Return or other electrode(s) | Duration per session (min) | Total sessions | Amplitude (as reported) |
| (Cavendish et al., 2022) | 119 (89/30) | Intervention (Retrieval practice) | 30 | NR | NR | Adult | Declarative | Recall | Accuracy | tDCS | BP-tDCS | F3 | FP2 | 20 | 1 | 2 |
| | | Intervention (Restudy) | 30 | NR | NR | | | | | tDCS | BP-tDCS | F3 | FP2 | 20 | 1 | 2 |
| | | Control (RP) | 29 | NR | NR | | | | | Sham | | F3 | FP2 | | 1 | |
| | | Cotrol (RS) | 30 | NR | NR | | | | | Sham | | F3 | FP2 | | 1 | |
| (Grover et al., 2022) | 120 (56/64) | Exp 1: DLPFC gamma | 20 | 9 | 11 | Aged | Declarative | Recall | Accuracy | tACS | HD-tACS | AF3 | Fp1, Fz, F1, FC3, F5, AF7, AFz, EX17 | 20 | 4 | 1.58 |
| | | Exp 2: DLPFC theta | 20 | 9 | 11 | | | | | tACS | HD-tACS | AF3 | Fp1, Fz, F1, FC3, F5, AF7, AFz, EX17 | 20 | 4 | 1.58 |
| | | Exp 2: IPL gamma | 20 | 10 | 10 | | | | | tACS | HD-tACS | CP5 | C3, T7, CP1, FC5, FT7, TP7, PO7, EX13 | 20 | 4 | 1.58 |
| | | Exp 1：IPL theta | 20 | 8 | 12 | | | | | tACS | HD-tACS | CP5 | C3, T7, CP1, FC5, FT7, TP7, PO7, EX13 | 20 | 4 | 1.58 |
| | | Exp1: control | 20 | 9 | 11 | | | | | Sham | | | | | 4 | |
| | | Exp2: control | 20 | 11 | 9 | | | | | Sham | | | | | 4 | |
| (Smits et al., 2023) | 72 (4/68) | Intervention | 37 | 2 | 35 | Adult | Working | Recognition | Accuracy | tDCS | BP-tDCS | F4 | C2 | 20 | 3 | 2 |
| | | Control | 35 | 2 | 33 | | | | | Sham | | F4 | C2 | | 3 | |
| (Friehs et al., 2021) | 78 (-/-) | Anodal | 26 | NR | NR | Adult | Declarative | Recognition | Accuracy | tDCS | MP-tDCS | BA 38 at the midpoint between F7, T7 and FT9 position | left deltoid | 20 | 1 | 0.7 |
| | | Cathodal | 26 | NR | NR | | | | | tDCS | MP-tDCS | left deltoid | BA 38 at the midpoint between F7, T7 and FT9 position | 20 | 1 | 0.7 |
| | | Control | 26 | NR | NR | | | | | Sham | | BA 38 at the midpoint between F7, T7 and FT9 position | left deltoid | | 1 | |
| (Bolling et al., 2021) | 58 (-/-) | Intervention (weak), A day after | 21 | NR | NR | Adult | Declarative | Recall | Accuracy | tDCS | BP-tDCS | F3 | F4 | 20 | 2 | 1 |
| | | Intervention (strong), A day after | 17 | NR | NR | | | | | tDCS | BP-tDCS | F3 | F4 | 20 | 2 | 1.5 |
| | | Control (Sham, A day after) | 20 | NR | NR | | | | | Sham | | F3 | F4 | | 2 | |
| (Assecondi et al., 2021) | 84 (-/-) | Low BS (with strategy) | 7 | NR | NR | Adult | Working | Recognition | Accuracy | tDCS | BP-tDCS | F4 | Fp1 | 20 | 2 | 2 |
| | | Low BS (witout strategy) | 9 | NR | NR | | | | | tDCS | BP-tDCS | F4 | Fp1 | 20 | 2 | 2 |
| | | Mid BS (with) | 6 | NR | NR | | | | | tDCS | BP-tDCS | F4 | Fp1 | 20 | 2 | 2 |
| | | Mid BS (without | 6 | NR | NR | | | | | tDCS | BP-tDCS | F4 | Fp1 | 20 | 2 | 2 |
| | | High BS (with) | 8 | NR | NR | | | | | tDCS | BP-tDCS | F4 | Fp1 | 20 | 2 | 2 |
| | | High BS (without) | 6 | NR | NR | | | | | tDCS | BP-tDCS | F4 | Fp1 | 20 | 2 | 2 |
| | | Control for Low BS (with strategy) | 6 | NR | NR | | | | | Sham | | F4 | Fp1 | | 2 | |
| | | Control for Low BS (witout strategy) | 6 | NR | NR | | | | | Sham | | F4 | Fp1 | | 2 | |
| | | Control for Mid BS (with) | 9 | NR | NR | | | | | Sham | | F4 | Fp1 | | 2 | |
| | | Control for Mid BS (without | 6 | NR | NR | | | | | Sham | | F4 | Fp1 | | 2 | |

| Study and participants | | | | | | Memory outcomes | | | | Brain stimulation | | | | | | |
|---|---|---|---|---|---|---|---|---|---|---|---|---|---|---|---|---|
| Reference | Total N (female/male) | Group | Group N | Female | Male | Age group | Memory type | Recall or recognition | Outcome | NIBS type | Montage | Anode (tDCS) or electrode 1 | Return or other electrode(s) | Duration per session (min) | Total sessions | Amplitude (as reported) |
| | | Control for High BS (with) | 6 | NR | NR | | | | | Sham | | F4 | Fp1 | | 2 | |
| | | Control for High BS (without) | 9 | NR | NR | | | | | Sham | | F4 | Fp1 | | 2 | |
| (Marko & Riečanský, 2021) | 121 (72/49) | Intervention (PFC) | 40 | NR | NR | Adult | Working | Recall | Accuracy, RT | tDCS | BP-tDCS | b/w F3 and AF3 | b/w T7 and P5 | 25 | 1 | 2 |
| | | Intervention (TPC) | 40 | NR | NR | | | | | tDCS | BP-tDCS | b/w T8 and P6 | b/w T7 and P5 | 25 | 1 | 2 |
| | | Control | 41 | NR | NR | | | | | Sham | | b/w F3 and AF3 or b/w T8 and P6 (balanced across participants) | b/w T7 and P5 | | 1 | |
| (Murphy et al., 2020) | 49 (33/16) | tDCS | 16 | 11 | 5 | Adult | Working | Recognition | Accuracy, RT | tDCS | BP-tDCS | F3 | right supraorbital | 20 | 1 | 1 |
| | | tRNS | 16 | 10 | 6 | | | | | tRNS | BP-tRNS | F3 | right supraorbital | 20 | 1 | 1 |
| | | Control | 17 | 12 | 5 | | | | | Sham | | F3 | right supraorbital | | 1 | |
| (Nissim et al., 2019) | 28 (15/13) | Intervention with training | 14 | 7 | 7 | Aged | Working | Recognition | Accuracy, RT | tDCS | BP-tDCS | F4 | F3 | 20 | 10 | 2 |
| | | Control with training | 14 | 8 | 6 | | | | | Sham | | F4 | F3 | | 10 | |
| (Lerner et al., 2019) | 24 (14/10) | Intervention | 12 | NR | NR | Adult | Declarative | Recall | Accuracy | tDCS, tACS | BP-tDCS + BP-tACS | tDCS: 32 stimulation electrodes located in the 64-channel 10/10 layout / tACS:in-phase F5 and F6 in the 10/10 system, and out-of-phase P5 and P6 at 1.5 mA/hemisphere | | | | tDCS: <=2.5; tACS: 3 (peak-to-peak) |
| | | Control | 12 | NR | NR | | | | | Sham | | | | | | |
| (Marián et al., 2018) | 118 (74/44) | Exp 1. Intervention | 33 | 23 | 10 | Adult | Declarative | Recall | Accuracy | tDCS | BP-tDCS | F4 | Cz | 15 | 1 | 2 |
| | | Exp 2. Intervention | 27 | 17 | 10 | | | | | tDCS | BP-tDCS | F4 | Cz | 15 | 1 | 2 |
| | | Exp 1. Control | 33 | 19 | 14 | | | | | Sham | | F4 | Cz | | 1 | |
| | | Exp 2. Control | 25 | 15 | 10 | | | | | Sham | | F4 | Cz | | 1 | |
| (Balzarotti & Colombo, 2016) | 42 (-/-) | Anodal | 14 | NR | NR | Adult | Declarative | Recall | Accuracy | tDCS | BP-tDCS | F3 | left mastoid | 15 | 1 | 1.5 |
| | | Cathodal | 14 | NR | NR | | | | | tDCS | BP-tDCS | left mastoid | F3 | 15 | 1 | 1.5 |
| | | Control | 14 | NR | NR | | | | | Sham | | F3 | left mastoid | | 1 | |
| (Chander et al., 2016) | 20 (4/16) | Intervention | 10 | NR | NR | Adult | Working | Recognition | Accuracy | tACS | BP-tACS | Fpz | Pz | 4 | 1 | 2 |
| | | Control | 10 | NR | NR | | | | | Sham | | Fpz | Pz | | 1 | |
| (Vosskuhl et al., 2015) | 33 (14/19) | Intervention | 17 | 7 | 10 | Adult | Working | Recall, Recognition | Accuracy | tACS | BP-tACS | FCz | Pz | 18 | 1 | 0.843 |
| | | Sham control | 16 | 7 | 9 | | | | | Sham | | FCz | Pz | | 1 | |
| (Martin et al., 2013) | 52 (22/30) | Intervention tDCS only | 10 | 5 | 5 | Adult | Working | Recall | Accuracy | tDCS | MP-tDCS | F3 | right deltoid | 30 | 10 | 2 |
| | | Intervention tDCS + CT | 21 | 9 | 12 | | | | | tDCS | MP-tDCS | F3 | right deltoid | 30 | 10 | 2 |
| | | Sham tDCS + CT | 21 | 8 | 13 | | | | | Sham | | F3 | right deltoid | | 10 | |
| (Smet et al., 2024) | 146 (107/39) | Intervention tDCS + control task | 36 | 26 | 10 | Adult | Working | Recognition | Accuracy, RT | tDCS | BP-tDCS | F3 | F4 | 30 | 1 | 2 |
| | | Intervention tDCS + stress task | 37 | 28 | 9 | | | | | tDCS | BP-tDCS | F3 | F4 | 30 | 1 | 2 |
| | | Sham tDCS + control task | 37 | 27 | 10 | | | | | Sham | | F3 | F4 | | 1 | |
| | | Sham tDCS + stress task | 36 | 26 | 10 | | | | | Sham | | F3 | F4 | | 1 | |
| (Gnedykh et al., 2024) | 72 (56/16) | Anodal | 24 | NR | NR | Adult | Declarative | Recognition | Accuracy, RT | tDCS | MP-tDCS | CP6 | rhomboid major muscle area along the spine on the same side as the anode (right) | 15 | 1 | 1.5 |
| | | Cathodal | 24 | NR | NR | | | | | tDCS | MP-tDCS | rhomboid major muscle area along the spine on the same side as the anode (right) | CP6 | 15 | 1 | 1.5 |

| Study and participants | | | | | | Memory outcomes | | | | Brain stimulation | | | | | | |
|---|---|---|---|---|---|---|---|---|---|---|---|---|---|---|---|---|
| Reference | Total N (female/male) | Group | Group N | Female | Male | Age group | Memory type | Recall or recognition | Outcome | NIBS type | Montage | Anode (tDCS) or electrode 1 | Return or other electrode(s) | Duration per session (min) | Total sessions | Amplitude (as reported) |
| | | Control | 24 | NR | NR | | | | | Sham | | CP6 | rhomboid major muscle area along the spine on the same side as the anode (right) | | 1 | |
| (Otstavnov et al., 2024) | 47 (24/23) | Double stim | 16 | NR | NR | Adult | Working | Recall | Accuracy, RT | HD-tDCS | HD-tDCS | F3 & P3 | F1, FC3, F5, AF3, CP3, P1, P5, PO3 | 15 | 1 | 1 |
| | | Single stim | 16 | NR | NR | | | | | HD-tDCS | HD-tDCS | F3 | F1, FC3, F5, AF3 | 15 | 1 | 1 |
| | | Control | 15 | NR | NR | | | | | Sham | | F3 & P3 | F1, FC3, F5, AF3, CP3, P1, P5, PO3 | | 1 | |
| (Hooyman et al., 2024) | 60 (23/37) | Intervention | 29 | 11 | 18 | Adult | Working | Recall | Accuracy, RT | tDCS | BP-tDCS | P4 | left supraorbital | 20 | 1 | 2 |
| | | Control | 31 | 12 | 19 | | | | | Sham | | P4 | left supraorbital | | 1 | |
| (Haciahmet et al., 2024) | 60 (48/11; 1 diverse) | Intervention | 30 | 23 | 7 | Adult | Declarative | Recall | Accuracy | tDCS | MP-tDCS | CP5 | left deltoid | 10 | 1 | 0.5 |
| | | Control | 30 | 25 | 4 | | | | | Sham | | CP5 | left deltoid | | 1 | |
| (Fromm et al., 2024) | 52 (40/12) | Intervention | 26 | 20 | 6 | Adult | Declarative | Recall | Accuracy | tDCS | HD-tDCS | CP6 | FC4 & FT8 & P2 & PO8 | 20 | 1 | 2 |
| | | Control | 26 | 20 | 6 | | | | | Sham | | CP6 | FC4 & FT8 & P2 & PO8 | | 1 | |
| (Edgcumbe et al., 2024) | 48 (27/21) | Anodal-anodal | 16 | 10 | 6 | Adult | Working | Recognition | Accuracy | tDCS | BP-tDCS | F4 | F3 | 20 | 2 | 1.5 |
| | | Anodal-Sham | 16 | 9 | 7 | | | | | tDCS + Sham | BP-tDCS | F4 | F3 | 20 | 2 | 1.5 |
| | | Control | 16 | 8 | 8 | | | | | Sham | | F4 | F3 | | 1 | |
| (Lezama et al., 2024) | 104 (44/60) | Exp1 Anodal | 32 | NR | NR | Adult | Declarative | Recall | Accuracy | tDCS | MP-tDCS | FT9 | right deltoid | 20 | 1 | 1.5 |
| | | Exp2 Cathodal | 20 | NR | NR | | | | | tDCS | MP-tDCS | left deltoid | F4 | 20 | 1 | 1.5 |
| | | Exp1 Control | 32 | NR | NR | | | | | Sham | | FT9 | right deltoid | | 1 | |
| | | Exp2 Control | 20 | NR | NR | | | | | Sham | | left deltoid | F4 | | 1 | |
| (Diedrich et al., 2024) | 35 (19/16) | Intervention | 17 | 10 | 7 | Aged | Working | Recognition | RT | tACS | BP-tACS | F3 | F4 | 20 | 16 | 2 |
| | | Control | 18 | 9 | 9 | | | | | Sham | | F3 | F4 | | 16 | |
| (Civile et al., 2024) | 120 (102/18) | Fp3-Fp2 | 40 | NR | NR | Adult | Working | Recognition | Accuracy | tDCS | BP-tDCS | Fp3 | Fp2 | 10 | 1 | 1.5 |
| | | Fp3-Cz | 40 | NR | NR | | | | | tDCS | BP-tDCS | Fp3 | Cz | 10 | 1 | 1.5 |
| | | Control | 40 | NR | NR | | | | | Sham | | Fp3 | Fp2 or Cz | | 1 | |
| (Öksüz & Eskikurt, 2024) | 42 (-/-) | Intervention | 21 | NR | NR | Adult | Working | Recognition | Accuracy | tDCS | BP-tDCS | left supraorbital | F3 | 20 | 1 | 2 |
| | | Control | 21 | NR | NR | | | | | Sham | | left supraorbital | F3 | | 1 | |
| (Haslacher et al., 2024) | 46 (19/27) | Parieto-occipital CLAM-tACS | 21 | 8 | 13 | Adult | Working | Recognition | Accuracy | tACS | MP-tACS | Oz | Cz | 45 | 1 | 20 |
| | | Frontal Intervention control | 25 | 11 | 14 | | | | | tACS | MP-tACS | Fpz | Cz | 45 | 1 | 20 |
| (Sandrini et al., 2024) | 80 (57/23) | Encoding - Anodal | 14 | NR | NR | Aged | Declarative | Recall | Accuracy | tDCS | BP-tDCS | F3 | right supraorbital | 15 | 1 | 1.5 |
| | | Consolidation - Anodal | 14 | NR | NR | | | | | tDCS | BP-tDCS | F3 | right supraorbital | 15 | 1 | 1.5 |
| | | Reconsolidation - Anodal | 12 | NR | NR | | | | | tDCS | BP-tDCS | F3 | right supraorbital | 15 | 1 | 1.5 |
| | | Encoding - Sham | 14 | NR | NR | | | | | Sham | | F3 | right supraorbital | | 1 | |
| | | Consolidation - Sham | 14 | NR | NR | | | | | Sham | | F3 | right supraorbital | | 1 | |
| | | Reconsolidation - Sham | 12 | NR | NR | | | | | Sham | | F3 | right supraorbital | | 1 | |
| (Riemann et al., 2024) | 156 (99/57) | IFG stim during acquisition | 27 | 17 | 10 | Adult | Declarative | Recall | Accuracy | tDCS | HD-tDCS | F7 | Fp1, F3, FC5 | 20 | 1 | 1.5 |
| | | MTG stim during acquisition | 27 | 23 | 4 | | | | | tDCS | HD-tDCS | TP7 | FT7, C5, P9, PO7 | 20 | 1 | 1.5 |
| | | IFG stim during recall | 25 | 14 | 11 | | | | | tDCS | HD-tDCS | F7 | Fp1, F3, FC5 | 20 | 1 | 1.5 |
| | | MTG stim during recall | 25 | 16 | 9 | | | | | tDCS | HD-tDCS | TP7 | FT7, C5, P9, PO7 | 20 | 1 | 1.5 |

| Study and participants | | | | | | Memory outcomes | | | | Brain stimulation | | | | | | |
|---|---|---|---|---|---|---|---|---|---|---|---|---|---|---|---|---|
| Reference | Total N (female/male) | Group | Group N | Female | Male | Age group | Memory type | Recall or recognition | Outcome | NIBS type | Montage | Anode (tDCS) or electrode 1 | Return or other electrode(s) | Duration per session (min) | Total sessions | Amplitude (as reported) |
| | | Sham during acquisition (50% IFG/MTG montage) | 26 | 11 | 15 | | | | | Sham | | F7 / TP7 (50% each) | Fp1, F3, FC5 / FT7, C5, P9, PO7 (50% each) | | 1 | |
| | | Sham during recall (50% IFG/MTG montage) | 26 | 18 | 8 | | | | | Sham | | F7 / TP7 (50% each) | Fp1, F3, FC5 / FT7, C5, P9, PO7 (50% each) | | 1 | |
| (Almeida et al., 2023) | 42 (-/-) | left DLPFC | 14 | NR | NR | Aged | Declarative | Recall | Accuracy | tDCS | MP-tDCS | F3 | right deltoid | 20 | 12 | 2 |
| | | Right cerebellum | 14 | NR | NR | | | | | tDCS | MP-tDCS | 1 cm under and 4 cm lateral to the inion | right deltoid | 20 | 12 | 2 |
| | | Control | 14 | NR | NR | | | | | Sham | | F3 | right deltoid | | 12 | |
| (Berglund-Barraza et al., 2023) | 28 (19/8; nonbinary=1) | Intervention | 14 | 9 | 5 | Adult | Working | Recall | Accuracy | HD-tDCS | HD-tDCS | Fz | Fpz & F7 & F8 & Cz | 20 | 1 | 1 |
| | | Control | 14 | 10 | 4 | | | | | Sham | | Fz | Fpz & F7 & F8 & Cz | | 1 | |
| (Ankri et al., 2023) | 130 (130/0) | Stress + Intervention tDCS | 33 | 33 | 0 | Adult | Working, Declarative | Recall, Recognition | Accuracy | tDCS | BP-tDCS | between F4 & AF4 | Cz | 20 | 1 | 2 |
| | | No stress + Intervention tDCS | 30 | 30 | 0 | | | | | tDCS | BP-tDCS | between F4 & AF4 | Cz | 20 | 1 | 2 |
| | | Stress + Sham | 34 | 34 | 0 | | | | | Sham | | between F4 & AF4 | Cz | | 1 | |
| | | No stress + Sham | 33 | 33 | 0 | | | | | Sham | | between F4 & AF4 | Cz | | 1 | |
| (Ke et al., 2023) | 34 (16/18) | Intervention | 17 | NR | NR | Adult | Working | Recognition | Accuracy, RT | HD-tDCS | HD-tDCS | F3 | Fp1 & Fz & C3 & F7 | 25 | 10 | 2 |
| | | Control | 17 | NR | NR | | | | | Sham | | F3 | Fp1 & Fz & C3 & F7 | | 10 | |
| (Bartl et al., 2023) | 80 (-/-) | Experiment 1 – Intervention tDCS | 19 | NR | NR | Adult | Declarative | Recall, Recognition | Accuracy | tDCS | BP-tDCS | F3 | Fp2 | 16 | 1 | 1.5 |
| | | Experiment 2 – Intervention tDCS | 20 | NR | NR | | | | | tDCS | BP-tDCS | F3 | Fp2 | 16 | 1 | 1.5 |
| | | Experiment 1 – Sham | 21 | NR | NR | | | | | Sham | | F3 | Fp2 | | 1 | |
| | | Experiment 2 – Sham | 22 | NR | NR | | | | | Sham | | F3 | Fp2 | | 1 | |
| (Balboa-Bandeira et al., 2023) | 64 (49/15) | tDCS | 16 | 12 | 4 | Adult | Declarative | Recall, Recognition | Accuracy | tDCS | BP-tDCS | FC5 | P5 | 20 | 1 | 1.5 |
| | | tRNS | 16 | 13 | 3 | | | | | tRNS | BP-tRNS | FC5 | P5 | 20 | 1 | 1.5 |
| | | tDCS + tRNS | 16 | 11 | 5 | | | | | tDCS + tRNS | BP-tDCS + BP-tRNS | FC5 | P5 | 20 | 1 | 1.5 |
| | | Sham | 16 | 13 | 3 | | | | | Sham | | FC5 | P5 | | 1 | |
| (Ma & Pang, 2023) | 128 (70/58) | Social inclusion + Intervention tDCS | 32 | NR | NR | Adult | Working | Recognition | Accuracy | tDCS | BP-tDCS | F6 | Fp1 | 20 | 1 | 2 |
| | | Social exclusion + Intervention tDCS | 32 | NR | NR | | | | | tDCS | BP-tDCS | F6 | Fp1 | 20 | 1 | 2 |
| | | Social inclusion + Sham | 32 | NR | NR | | | | | Sham | | F6 | Fp1 | | 1 | |
| | | Social exclusion + Sham | 32 | NR | NR | | | | | Sham | | F6 | Fp1 | | 1 | |
| (Figeys et al., 2023) | 32 (20/12) | Intervention | 16 | 10 | 6 | Adult | Working | Recognition | Accuracy, RT | tDCS | MP-tDCS | F4 | left deltoid | 20 | 1 | 1.5 |
| | | Control | 16 | 10 | 6 | | | | | Sham | | F4 | left deltoid | | 1 | |
| (Maldonado et al., 2023) | 72 (-/-) | Anodal | 23 | NR | NR | Adult | Working | Recognition | Accuracy, RT | tDCS | MP-tDCS | 2 cm below and 4 cm right-lateral to the inion (right cerebellum) | right deltoid | 20 | 1 | 2 |
| | | Cathodal | 25 | NR | NR | | | | | tDCS | MP-tDCS | right deltoid | 2 cm below and 4 cm right-lateral to the inion (right cerebellum) | 20 | 1 | 2 |

| Study and participants | | | | | | Memory outcomes | | | | Brain stimulation | | | | | | |
|---|---|---|---|---|---|---|---|---|---|---|---|---|---|---|---|---|
| Reference | Total N (female/male) | Group | Group N | Female | Male | Age group | Memory type | Recall or recognition | Outcome | NIBS type | Montage | Anode (tDCS) or electrode 1 | Return or other electrode(s) | Duration per session (min) | Total sessions | Amplitude (as reported) |
| | | Sham | 24 | NR | NR | | | | | Sham | | 2 cm below and 4 cm right-lateral to the inion (right cerebellum) | right deltoid | | 1 | |
| (Ji et al., 2023) | 60 (38/22) | tDCS + aerobic exercise | 20 | 12 | 8 | Adult | Working | Recall, Recognition | Accuracy | tDCS | BP-tDCS | F3 | Fp2 | 30 | 20 | 2 |
| | | tDCS only | 20 | 13 | 7 | | | | | tDCS | BP-tDCS | F3 | Fp2 | 30 | 20 | 2 |
| | | Aerobic exercise + Sham tDCS | 20 | 13 | 7 | | | | | Sham | | F3 | Fp2 | | 20 | |
| (Han et al., 2023) | 63 (-/-) | Left-anode | 21 | NR | NR | Adult | Declarative | Recognition | Accuracy | HD-tDCS | HD-tDCS | F5 | C1, C5, F9, Fp1 | 20 | 1 | 2 |
| | | Right-anode | 20 | NR | NR | | | | | HD-tDCS | HD-tDCS | F6 | C2, C6, F10, Fp2 | 20 | 1 | 2 |
| | | Combined Sham (left/right montage, 11 each) | 22 | NR | NR | | | | | Sham | | F5 / F6 | C1, C5, F9, Fp1 / C2, C6, F10, Fp2 (11 each) | | 1 | |
| (Moghadas Tabrizi et al., 2023) | 54 (27/27) | F4 Intervention | 14 | NR | NR | Adult | Working | Recall | Accuracy, RT | tDCS | BP-tDCS | F4 | left supraorbital | 15 | 5 | 1.5 |
| | | P4 Intervention | 13 | NR | NR | | | | | tDCS | BP-tDCS | P4 | left supraorbital | 15 | 5 | 1.5 |
| | | F4 Sham | 13 | NR | NR | | | | | Sham | | F4 | left supraorbital | | 5 | |
| | | P4 Sham | 14 | NR | NR | | | | | Sham | | P4 | left supraorbital | | 5 | |
| (Zhang et al., 2022) | 28 (15/13) | Intervention | 18 | 9 | 9 | Adult | Working | Recognition | Accuracy, RT | tACS | BP-tACS | F4 & P4 | T8 | 15 | 1 | 2 |
| | | Control | 10 | 6 | 4 | | | | | Sham | | F4 & P4 | T8 | | 1 | |
| (Arar et al., 2024) | 193 (121/72) | Adults – left dlPFC Intervention | 51 | NR | NR | Adult | Working, Declarative | Recognition | Accuracy | tDCS | BP-tDCS | F3 | contralateral supraorbital | 20 | 2 | 2 |
| | | Older adults – left dlPFC Intervention | 46 | NR | NR | Aged | | | | tDCS | BP-tDCS | F3 | contralateral supraorbital | 20 | 2 | 2 |
| | | Adults – Sham | 51 | NR | NR | Adult | | | | Sham | | F3 | contralateral supraorbital | | 2 | |
| | | Older adults – Sham | 45 | NR | NR | Aged | | | | Sham | | F3 | contralateral supraorbital | | 2 | |
| (Beiman et al., 2025) | 30 (17/13) | Typically developed controls – Intervention tDCS | 15 | NR | NR | Adult | Working | Recognition | Accuracy | tDCS | BP-tDCS | F3 | F4 | 20 | 1 | 2 |
| | | Typically developed controls – Sham | 15 | NR | NR | | | | | Sham | | F3 | F4 | | 1 | |
| (Ben Izhak et al., 2025) | 58 (46/12) | Cognitive training + Intervention tDCS | 28 | 24 | 4 | Aged | Declarative | Recall | Accuracy | tDCS | BP-tDCS | F3 | above right eyebrow | 15 | 32 | 1 |
| | | Cognitive training + Sham tDCS | 30 | 22 | 8 | | | | | Sham | | F3 | above right eyebrow | 15 | 32 | |
| (Civile et al., 2025) | 80 (-/-) | Experiment 3: Sham → cathodal | 40 | NR | NR | Adult | Declarative | Recognition | Accuracy | tDCS | BP-tDCS | Fp3 | Fp2 | 10 | 2 | 1.5 |
| | | Experiment 3: Sham → Sham | 40 | NR | NR | | | | | Sham | | Fp3 | Fp2 | | 2 | |
| (Diedrich et al., 2025) | 77 (45/32) | Intervention | 38 | 21 | 17 | Aged | Declarative | Recall | Accuracy | tACS | BP-tACS | F3 | F4 | 20 | 16 | 2 |
| | | Control | 39 | 24 | 15 | | | | | Sham | | F3 | F4 | | 16 | |
| (Juras et al., 2025) | 58 (-/-) | Intervention | 29 | NR | NR | Adult | Declarative | Recall, Recognition | Accuracy | tDCS | MP-tDCS | P3 | right cheek | 20 | 3 | 1.5 |
| | | Control | 29 | NR | NR | | | | | Sham | | P3 | right cheek | | 3 | |
| (Khanmohammadi et al., 2025) | 80 (69/11) | right Cerebellum | 20 | 18 | 2 | Aged | Working | Recall | Accuracy | tDCS | HD-tDCS | Iz | Oz, O2, P8, PO8 | 20 | 1 | 2 |
| | | M1 | 20 | 17 | 3 | | | | | tDCS | HD-tDCS | Cz | Fz, C3, C4, Pz | 20 | 1 | 2 |
| | | left DLPFC | 20 | 18 | 2 | | | | | tDCS | HD-tDCS | F3 | Fz, C3, FP1, F7 | 20 | 1 | 2 |

| Study and participants | | | | | | Memory outcomes | | | | Brain stimulation | | | | | | |
|---|---|---|---|---|---|---|---|---|---|---|---|---|---|---|---|---|
| Reference | Total N (female/male) | Group | Group N | Female | Male | Age group | Memory type | Recall or recognition | Outcome | NIBS type | Montage | Anode (tDCS) or electrode 1 | Return or other electrode(s) | Duration per session (min) | Total sessions | Amplitude (as reported) |
| | | Control | 20 | 16 | 4 | | | | | Sham | | Iz / Cz / F3 (random) | Oz, O2, P8, PO8 / Fz, C3, C4, Pz / Fz, C3, Fp1, F7 (corresponding; random) | | 1 | |
| (Kianimotlagh et al., 2025) | 45 (-/-) | tDCS | 15 | NR | NR | Aged | Working | Recognition | Accuracy, RT | tDCS | BP-tDCS | F3 | F4 | 20 | 10 | 2 |
| | | tDCS+AVG | 15 | NR | NR | | | | | tDCS | BP-tDCS | F3 | F4 | 20 | 5 | 2 |
| | | Control | 15 | NR | NR | | | | | No stimulation | | | | | | |
| (Kukuła et al., 2025) | 90 (45/45) | Intervention | 30 | 15 | 15 | Adult | Working, Declarative | Recall | Accuracy | tDCS | BP-tDCS | F3 | FP2 | 30 | 10 | 2 |
| | | Control | 30 | 15 | 15 | | | | | Sham | BP-tDCS | F3 | FP2 | | 10 | |
| | | Control (no stimulation) | 30 | 15 | 15 | | | | | No stimulation | | | | | | |
| (Lin et al., 2025) | 54 (-/-) | PFC-Intervention | 18 | NR | NR | Adult | Working | Recall | Accuracy, RT | tDCS | MP-tDCS | F4 | left cheek | 20 | 1 | 2 |
| | | PPC-Intervention | 19 | NR | NR | | | | | tDCS | MP-tDCS | P4 | left cheek | 20 | 1 | 2 |
| | | PFC-Control | 18 | NR | NR | | | | | Sham | | F4 | left cheek | | 1 | |
| | | PPC-Control | 19 | NR | NR | | | | | Sham | | P4 | left cheek | | 1 | |
| (P. Lu & Yu, 2025) | 24 (12/12) | Intervention | 12 | 6 | 6 | Adult | Working | Recognition | Accuracy, RT | tDCS | BP-tDCS | F3 | FP2 | 20 | 1 | 1.5 |
| | | Control | 12 | 6 | 6 | | | | | Sham | | F3 | FP2 | | 1 | |
| (Mansur et al., 2025) | 51 (16/35) | Theta | 17 | NR | NR | Adult | Working | Recognition | Accuracy | tACS | BP-tACS | AF3 | CP5 | 15 | 1 | 2 |
| | | Alpha | 17 | NR | NR | | | | | tACS | BP-tACS | AF3 | CP5 | 15 | 1 | 2 |
| | | Control | 17 | NR | NR | | | | | Sham | | AF3 | CP5 | | 1 | |
| (Pan et al., 2025) | 118 (-/-) | ReI_Intervention | 30 | NR | NR | Adult | Declarative | Recall | Accuracy | tDCS | BP-tDCS | Oz | PO3, PO4, PO9, PO10 | 10 | 1 | 1 |
| | | ReI_Control | 29 | NR | NR | | | | | Sham | | Oz | PO3, PO4, PO9, PO10 | | 1 | |
| | | Nor_Intervention | 29 | NR | NR | | | | | tDCS | BP-tDCS | Oz | PO3, PO4, PO9, PO10 | 10 | 1 | 1 |
| | | Nor_Control | 30 | NR | NR | | | | | Sham | | Oz | PO3, PO4, PO9, PO10 | | 1 | |
| (Rosato et al., 2025) | 33 (-/-) | Intervention | 17 | NR | NR | | Working | Recognition | Accuracy, RT | tACS | HD-tACS | F3 | F7, Fp1, Fz, C3 | 20 | 3 | 1.5 |
| | | Control | 16 | NR | NR | | | | | Sham | | F3 | F7, Fp1, Fz, C3 | | 3 | |
| (Roy et al., 2025) | 75 (0/75) | Anodal dlPFC | 25 | 0 | 25 | | Working | Recognition | Accuracy | tDCS | HD-tDCS | F3 | Fp1, Fz, C3, F7 | 17 | 2 | 2 |
| | | Anodal vmPFC | 24 | 0 | 24 | | | | | tDCS | HD-tDCS | Nz | F7, F8, Ex19, Ex20 | 17 | 2 | 2 |
| | | Control | 26 | 0 | 26 | | | | | Sham | | | | | 2 | |
| (Shtoots et al., 2025) | 175 (102/73) | Theta frontal | 25 | 18 | 7 | Adult | Declarative | Recognition | Accuracy | tACS | HD-tACS | Fz | AF4, AF3, FC2, FC1 | 20 | 1 | 1.5 |
| | | Beta frontal | 25 | 12 | 13 | | | | | tACS | HD-tACS | Fz | AF4, AF3, FC2, FC1 | 20 | 1 | 1.5 |
| | | Theta left parietal | 25 | 16 | 9 | | | | | tACS | HD-tACS | P3 | Pz, O1, TP7, C3 | 20 | 1 | 1.5 |
| | | Beta left parietal | 25 | 17 | 8 | | | | | tACS | HD-tACS | P3 | Pz, O1, TP7, C3 | 20 | 1 | 1.5 |
| | | Theta bilateral parietal | 25 | 13 | 12 | | | | | tACS | BP-tACS | P1, P4 | chin | 20 | 1 | 1.5 |
| | | Beta bilateral parietal | 25 | 11 | 14 | | | | | tACS | BP-tACS | P1, P4 | chin | 20 | 1 | 1.5 |
| | | Control | 25 | 15 | 10 | | | | | Sham | | Fz | AF4, AF3, FC2, FC1 | | 1 | |
| (Sun et al., 2025) | 28 (10/18) | Intervention | 14 | 6 | 8 | Adult | Declarative | Recall | Accuracy | tACS | MP-tACS | P3 | right deltoid muscle | 20 | 1 | 1.5 |
| | | Control | 14 | 4 | 10 | | | | | Sham | | P3 | right deltoid muscle | | 1 | |
| (Zacharopoulos et al., 2025) | 72 (36/36) | DLPFC Intervention | 24 | 12 | 12 | Adult | Working | Recall | Accuracy, RT | tRNS | BP-tRNS | F3 | F4 | 20 | 5 | 1 |
| | | PPC Intervention | 24 | 12 | 12 | | | | | tRNS | BP-tRNS | P3 | P4 | 20 | 5 | 1 |

| Study and participants | | | | | | Memory outcomes | | | | Brain stimulation | | | | | | |
|---|---|---|---|---|---|---|---|---|---|---|---|---|---|---|---|---|
| Reference | Total N (female/male ) | Group | Group N | Femal e | Ma le | Age group | Memory type | Recall or recognition | Outcom e | NIBS type | Montage | Anode (tDCS) or electrode 1 | Return or other electrode(s) | Duration per session (min) | Total sessions | Amplitud e (as reported) |
| | | Control | 24 | NR | NR | | | | | Sham | | F3 / P3 (half each) | F4 / P4 (corresponding; half each) | | 5 | |
| (Raviv et al., 2025) | 30 (18/12) | Intervention | 15 | 9 | 6 | Adult | Working | Recognition | Accurac y, RT | tACS | BP-tACS | F3 | F4 | 20 | 5 | 2 |
| | | Control | 15 | 9 | 6 | | | | | Sham | | F3 | F4 | | 5 | |
| (Liu et al., 2025) | 180 (105/75) | Exp1 anodal | 30 | 14 | 16 | Adult | Working | Recognition | Accurac y, RT | tDCS | HD-tDCS | CP6 | C4, P4, P8, T8 | 20 | 1 | 2 |
| | | Exp1 cathodal | 30 | 12 | 18 | | | | | tDCS | HD-tDCS | C4, P4, P8, T8 | CP6 | 20 | 1 | 2 |
| | | Exp2 anodal | 30 | 19 | 11 | | | | | tDCS | HD-tDCS | CP6 | C4, P4, P8, T8 | 20 | 1 | 2 |
| | | Exp2 cathodal | 30 | 22 | 8 | | | | | tDCS | HD-tDCS | C4, P4, P8, T8 | CP6 | 20 | 1 | 2 |
| | | Exp1 Control | 30 | 19 | 11 | | | | | Sham | | CP6 or C4, P4, P8, T8 (half each) | C4, P4, P8, T8 or CP6 (inverse; half each) | | 1 | |
| | | Exp2 Control | 30 | 19 | 11 | | | | | Sham | | CP6 or C4, P4, P8, T8 (half each) | C4, P4, P8, T8 or CP6 (inverse; half each) | | 1 | |
| (Bueno et al., 2025) | 30 (30/0) | Intervention | 10 | 10 | 0 | Aged | Declarativ e | Recall | Accurac y | tDCS | BP-tDCS | F3 | F7 | 20 | 9 | 2 |
| | | Control | 10 | 10 | 0 | | | | | Sham | | F3 | F7 | | 9 | |
| | | Control (CCT only) | 10 | 10 | 0 | | | | | No stimulation | | | | | | |
