## Supplementary Figures for "Transcranial electrical stimulation for memory enhancement: A systematic review and meta-analysis"

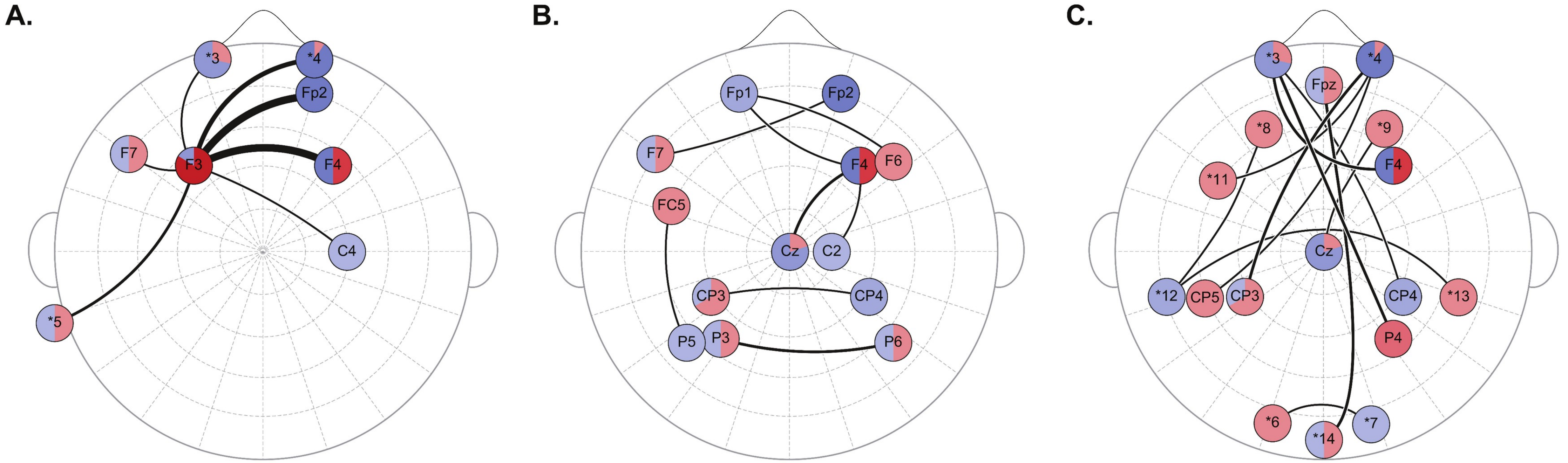


**Supplementary Figure 1.** Full bipolar tDCS electrode montage maps. All bipolar tDCS pairs with identifiable electrode locations are shown, including pairs with a summed count of one; Figure 3B shows the corresponding subset with summed counts of at least two. Pairs are grouped as (A) connections involving F3, (B) remaining connections between standard scalp electrode locations, and (C) remaining connections involving nonstandard or extracephalic locations. Connections involving Fp3 are omitted because its scalp location could not be established; no coordinate was inferred. Red and blue sectors indicate anodal and cathodal use, respectively, and their proportions reflect article counts summed across all bipolar tDCS configurations. An article reporting multiple configurations or polarities may contribute more than once. Colour intensity represents counts in four bands (1, 2–3, 4–7, and ≥8); circular nodes have a uniform size and do not encode counts. Black curves connect paired electrodes; line thickness represents the sum of the article counts for the two electrode directions. An article using both polarities can contribute to both directions. Asterisk labels shared with Figure 3 have the same definitions. Additional nonstandard placements are *11, the midpoint of the line joining (a) the intersection of the T7–F3 and F7–C3 lines and (b) the midpoint between F7 and F3; *12, the midpoint between T7 and P5; and *13, the midpoint between T8 and P6. T7 is the current designation for the former T3 position. tDCS, transcranial direct current stimulation.

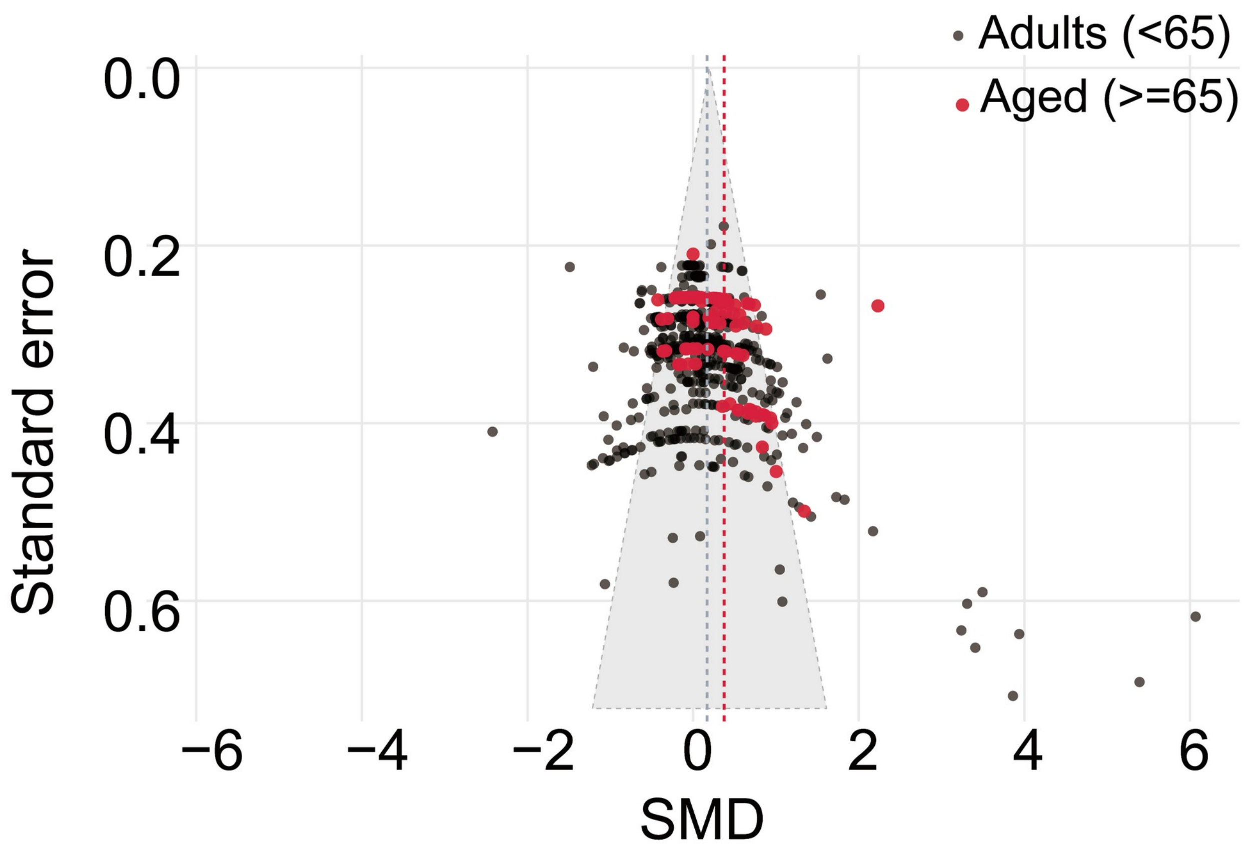


**Supplementary Figure 2.** Funnel plot of tDCS versus sham comparisons in younger and older adults. Each point is one effect size (Hedges' g; positive values indicate better memory performance with active stimulation) plotted against its standard error; the vertical axis is inverted so that more precise estimates lie at the top. Grey points show younger adults (<65 years; 483 effect sizes from 78 articles) and red points show older adults (≥65 years; 102 effect sizes from 15 articles), drawn on top. The solid vertical line and shaded triangle indicate the pooled tDCS estimate for both age groups combined (g = 0.20) and its pseudo 95% confidence limits (±1.96 standard errors). Dashed vertical lines indicate the subgroup estimates from the multilevel random-effects models (younger adults, g = 0.17 [95% CI 0.09–0.25]; older adults, g = 0.38 [0.19–0.56]). Effect sizes from the same article often share one standard error and therefore form horizontal bands. At the article level, older-adult studies used more stimulation days (median 7 vs 1; Wilcoxon rank-sum p = 0.0015) and a longer total stimulation time (median 140 vs 20 min; p = 0.0046). Furthermore, the older-adult subgroup also showed funnel-plot asymmetry (Egger's test Z = 3.65; Begg's rank correlation τ = 0.37, compared with Z = 13.73 and τ = 0.09 in younger adults). Which subgroup appears more asymmetric therefore depends on the test used, but publication bias remains a concern in this group.

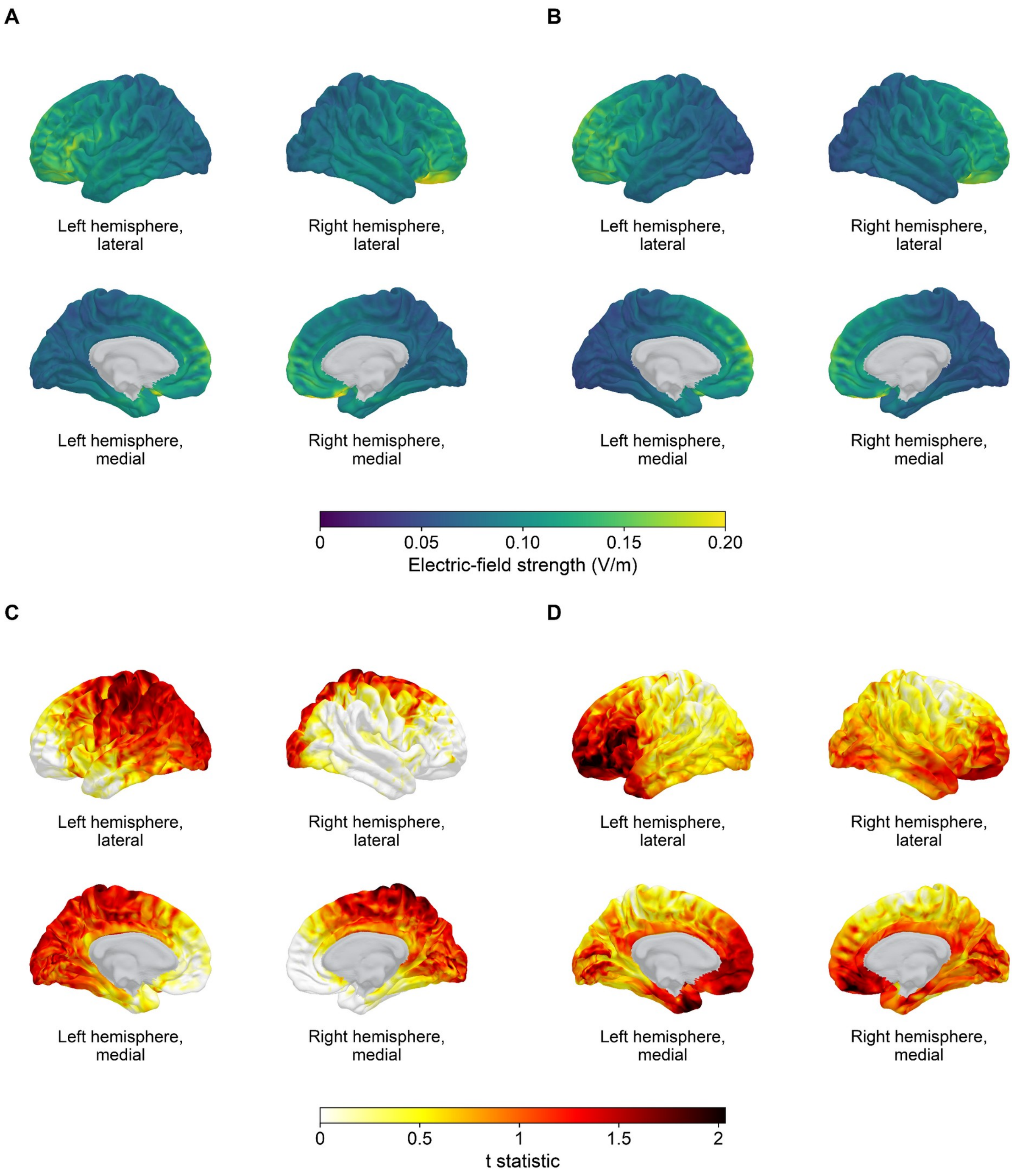


**Supplementary Figure 3.** Electric-field distributions and vertex-wise t-statistics in declarative- and working-memory studies. The left column shows declarative memory (DM) and the right column shows working memory (WM). (A, B) Mean electric-field strength at each fsaverage vertex across distinct study-montage pairs (DM: 51 pairs from 39 studies; WM: 57 pairs from 44 studies). Values above 0.2 V/m use the upper endpoint colour. (C, D) Vertex-wise t-statistics for associations between electric-field strength and Hedges' g. Models adjusted for age group and memory test timing relative to tDCS, with standard errors accounting for dependence among observations from the same study. Analyses included 60 aggregated observations for DM and 65 for WM. Positive t values are displayed regardless of statistical significance; nonpositive values are white. Grey indicates vertices outside the HCP-MMP analysis mask.
