## Supplementary Table 2 for "Transcranial electrical stimulation for memory enhancement: A systematic review and meta-analysis"

**Supplementary Table 2.** Pooled effects, heterogeneity, and small-study effects for the overall analysis and each subgroup.

*Positive standardised mean differences (SMD, Hedges’ g) indicate better memory performance with active than with sham stimulation. Bold: 95% CI excludes 0.*

| **Subgroup** | **Trials** | **Articles** | **Effect sizes (k)** | **Participants** **(active / sham)** | **SMD [95% CI]** | **Cochran’s Q** | **Egger’s test** | **Begg’s test** | **Risk of bias (%)** **low / unclear / high** |
| --- | --- | --- | --- | --- | --- | --- | --- | --- | --- |
| ***Overall*** | | | | | | | | | |
| All trials (all stimulation types) | 193 | 108 | 738 | 3,988 / 2,829 | **0.19 [0.12, 0.25]** | Q(737) = 1725.45, p < .001 | Z = 13.81, p < .001 | τ = 0.09, p < .001 | 40 / 60 / 0 |
| ***Stimulation type*** *(omnibus test of differences: QM(1) = 0.07, p = .788)* | | | | | | | | | |
| tDCS | 155 | 90 | 585 | 3,292 / 2,497 | **0.20 [0.12, 0.27]** | Q(584) = 1478.27, p < .001 | Z = 15.32, p < .001 | τ = 0.12, p < .001 | 40 / 60 / 0 |
| tACS | 30 | 16 | 103 | 560 / 296 | **0.18 [0.03, 0.33]** | Q(102) = 192.43, p < .001 | Z = −1.67, p = .095 | τ = −0.18, p = .007 | 38 / 62 / 0 |
| ***tDCS: electrode montage*** *(omnibus test of differences: QM(2) = 0.63, p = .731)* | | | | | | | | | |
| Monopolar | 45 | 25 | 183 | 906 / 595 | **0.20 [0.11, 0.29]** | Q(182) = 305.63, p < .001 | Z = 7.01, p < .001 | τ = 0.14, p = .004 | 28 / 72 / 0 |
| Bipolar | 83 | 50 | 317 | 1,783 / 1,539 | **0.22 [0.09, 0.35]** | Q(316) = 1000.48, p < .001 | Z = 13.28, p < .001 | τ = 0.06, p = .088 | 46 / 54 / 0 |
| High-definition (HD) | 26 | 14 | 79 | 578 / 338 | 0.14 [−0.03, 0.30] | Q(78) = 150.50, p < .001 | Z = 4.25, p < .001 | τ = 0.27, p < .001 | 43 / 57 / 0 |
| ***tDCS: age group*** *(omnibus test of differences: QM(1) = 17.75, p < .001)* | | | | | | | | | |
| Younger adults (<65 y) | 134 | 78 | 483 | 2,863 / 2,150 | **0.17 [0.09, 0.25]** | Q(482) = 1285.76, p < .001 | Z = 13.73, p < .001 | τ = 0.09, p = .004 | 35 / 65 / 0 |
| Older adults (≥65 y) | 21 | 15 | 102 | 429 / 347 | **0.38 [0.19, 0.56]** | Q(101) = 186.76, p < .001 | Z = 3.65, p < .001 | τ = 0.37, p < .001 | 67 / 33 / 0 |
| ***tDCS: memory type*** *(omnibus test of differences: QM(1) = 0.04, p = .842)* | | | | | | | | | |
| Working memory | 86 | 51 | 335 | 1,838 / 1,416 | **0.19 [0.09, 0.29]** | Q(334) = 868.65, p < .001 | Z = 12.82, p < .001 | τ = 0.06, p = .086 | 43 / 57 / 0 |
| Declarative memory | 74 | 43 | 223 | 1,694 / 1,335 | **0.21 [0.11, 0.32]** | Q(222) = 543.81, p < .001 | Z = 8.99, p < .001 | τ = 0.28, p < .001 | 42 / 58 / 0 |
| ***tDCS: timing of assessment*** *(omnibus test of differences: QM(4) = 3.13, p = .537)* | | | | | | | | | |
| During stimulation | 42 | 21 | 101 | 838 / 636 | **0.22 [0.04, 0.41]** | Q(100) = 286.45, p < .001 | Z = 8.42, p < .001 | τ = 0.05, p = .491 | 38 / 62 / 0 |
| Within a few hours | 91 | 58 | 307 | 1,977 / 1,509 | **0.15 [0.06, 0.24]** | Q(306) = 811.94, p < .001 | Z = 8.54, p < .001 | τ = 0.13, p = .001 | 45 / 55 / 0 |
| 1 day | 22 | 15 | 57 | 482 / 426 | **0.31 [0.06, 0.56]** | Q(56) = 158.16, p < .001 | Z = 6.89, p < .001 | τ = 0.39, p < .001 | 40 / 60 / 0 |
| 1 week | 12 | 8 | 38 | 276 / 242 | 0.32 [−0.04, 0.68] | Q(37) = 67.05, p = .002 | Z = 3.11, p = .002 | τ = 0.53, p < .001 | 38 / 62 / 0 |
| > 1 week | 16 | 12 | 82 | 330 / 264 | 0.14 [−0.01, 0.29] | Q(81) = 153.19, p < .001 | Z = 5.05, p < .001 | τ = −0.14, p = .065 | 50 / 50 / 0 |
| ***tDCS: stimulation site*** | | | | | | | | | |
| Frontal target (HD or monopolar) | 39 | 27 | 156 | 721 / 560 | **0.28 [0.16, 0.39]** | Q(155) = 298.18, p < .001 | Z = 9.11, p < .001 | τ = 0.17, p = .002 | 33 / 67 / 0 |

Abbreviations: CI, confidence interval; HD, high-definition; k, number of effect sizes (comparisons); SMD, standardised mean difference (Hedges’ g); tACS, transcranial alternating current stimulation; tDCS, transcranial direct current stimulation.

Effect sizes were pooled with a multilevel random-effects model (rma.mv; random intercepts for article, study group, and task). “Overall” includes all stimulation types (8 trials of tRNS or combined protocols are included only in this row). Subgroups other than stimulation type are restricted to tDCS.

Trials are distinct active arms; participants are summed over distinct active and sham arms, each counted once. Cochran’s Q tests residual heterogeneity.

Egger’s test: multilevel version, with the standard error entered as a moderator in the same model. Begg’s test: rank correlation test treating effect sizes as independent (approximate). A criterion of p < .1 was used for both tests.

Risk of bias: percentage of articles in each subgroup rated low, unclear (intermediate), or high overall by the operational scoring rule described in Methods 2.3. All subgroups were rated as direct (no indirectness downgrade).

Stimulation site: tDCS trials with HD or monopolar montages whose target was coded as frontal. Omnibus tests (QM) compare the levels of each subgroup dimension within the corresponding multilevel meta-regression model.
